# Replication stress at centromeres biases the segregation of DNA damage

**DOI:** 10.64898/2026.08.31.748202

**Authors:** Elena Di Tommaso, Luigi Fanelli, Simona Giunta

## Abstract

Replication-associated errors can cause DNA damage to accumulate on the newly synthesized strand over time. In specific cases such as stem cells, retention of the “immortal” strand used as template preserves one daughter cell into pluripotency while correlating with terminal differentiation of the damage one. In somatic cells, DNA damage distribution after mitosis remains unclear. Here, we uncovered a mechanism of non-random segregation of the DNA damage marker γH2AX occurring during a single cell division cycle. Replication stress using hydroxyurea (HU) upon release into S phase in RPE-1, BJ, hCEC D29 and fibroblasts showed reproducible Non-Random Segregation (NRS) of γH2AX in the ensuing G1, a phenotype not observed in any of the cancer cell lines analysed. Notably, removal of R-loops led to a reduction of cells with NRS, whether RNaseH1 was over-expressed globally or exclusively targeted to centromeres, indicating that centromeric DNA-RNA hybrids contribute to NRS of the damage. In line with our previous evidence of centromeric chromatin disruption leading to R-loops, rapid removal of the histone H3 variant CENP-A causes damage and NRS, although to a lower extent than HU alone. This implies that additional mechanisms contribute to centromeric R-loops and NRS of damage in the daughter cells upon mitotic exit. Mechanistically, chemical inhibition of the catalytic activity of Rad51 led to a significant drop in NRS without a change in the total amount of damaged cells, implying involvement of the Homologous Recombination (HR) pathway to accumulation of γH2AX to only one chromatid. In turn, this affects the spindle-kinetochore with a measurable length asymmetry, inducing mechanical and/or epigenetic signals that affect the orientation of the sister chromatids on the metaphase plate to bias segregation. Altogether, we found replication-induced asymmetric segregation of DNA damage during mitosis that is influenced by centromeric R-loops, Rad51 activity and spindle dynamics, with implications on cell fate, chromosome and genome stability in the daughter cells.

## INTRODUCION

Centromeres are epigenetically specified by the presence of the centromere-specific histone H3 variant, CENP-A, which enables kinetochore assembly for microtubule attachment (Schueler and Sullivan, 2006; Santaguida and Musacchio, 2009; McKinley and Cheeseman, 2016). The kinetochore monitors the correct bi-orientation of microtubules to ensure the proper segregation of the sister chromatids. In the presence of unattached or incorrectly attached chromosomes, the dynamic nature of the attachment allows multiple attempts to establish the correct bi-orientation before the binding is stabilized and segregation proceeds (Musacchio, 2015). To ensure the correct binding of kinetochore proteins for a faithful chromosome segregation, is important for centromeric chromatin to properly fold in order to expose CENP-A nucleosomes to the kinetochore interface. We recently proposed a centromeric DNA conformation named the “layered looping model” where centromeric chromatin is organized into concentric loops arranged in a ring-like structure and layered with the flanking pericentromere (Di Tommaso and Giunta, 2023; Di Tommaso et al., 2023). While direct evidence for how the centromeric DNA is folded to allow this interaction are lacking, our model seeks to reconcile several previously proposed models (Schalch and Steiner, 2017), the solenoid/repeat-subunit model, which proposes that the chromatin is coiled from the *p* arm to the *q* arm across the primary constriction (Zinkowski et al., 1991; Blower et al., 2002; Sullivan and Karpen, 2004; Birchler et al., 2009); the looping model, based on the repetition of the single kinetochore module of the budding yeast (Dalal et al., 2007; Yeh et al., 2008); and the layered boustrophedon, based on the observation that CENP-A and H3 are jointly exposed on the surface of the centromeric chromatin to interact with kinetochore proteins (Ribeiro et al., 2010). Our additional finding that the ring-like structure is retained even in absence of spindle microtubules suggests that the organization of an outer CENP-A layer is intrinsic to centromeric architecture, independent of tension (Di Tommaso and Giunta, 2023; Di Tommaso et al., 2023).

Centromere integrity and stability are crucial to ensure the correct chromosome segregation, preserving the stability of intra- and inter-generational inheritance of the genetic information. In humans, centromeric DNA is organized as tandem arrays of ∼171 bp alpha-satellite repeats, further arranged into higher-order repeat (HOR) units spanning several megabases (Willard, 1985; Willard and Waye, 1987). This repetitive nature makes centromeres susceptible to DNA damage (Black and Giunta, 2018), to complex secondary structures (Kasinathan and Henikoff, 2018; Chittoor and Giunta, 2024), and prone to recombination events (Giunta and Funabiki, 2017; Giunta et al., 2021), a mechanism important not only for centromere evolution, but also as a response to exogenously induced DNA damage (Yilmaz et al., 2021). Rearrangements and recombination events at these repetitive sequences can lead to whole arm events, translocations (Giunta et al., 2021) as well as unequal sister chromatid exchanges (SCEs) of alpha-satellite DNA, a phenomenon exacerbated in cancer cell lines displaying either high- or low-chromosomal instability (CIN), and in cells approaching replicative senescence (Giunta and Funabiki, 2017). The presence of centromeric R-loops (DNA-RNA hybrids) contributes to DNA repair by facilitating the initiation of Homologous Recombination (HR) during the G1 phase (Yilmaz et al., 2021). During the late S and G2 phases however, when centromeres are synthesized, their accumulation is deleterious to centromere stability, as they interfere with DNA replication completion, transcription-replication collisions (Giunta et al., 2021) and/or with double strand breaks (DSBs) repair (Marnef and Legube, 2021; Ortega et al., 2021). In this context, core centromere proteins, and more specifically CENP-A and its associated proteins, play pivotal roles in preventing excessive recombination, and maintaining centromere integrity in human cells through repression of R-loops accumulation (Giunta and Funabiki, 2017; Giunta et al., 2021). Consistent with this, DNA breaks have been recently reported using exonuclease FISH (Exo-FISH) in both cancerous and non-cancerous, non-proliferating cells, confirms an intrinsic degree of instability (Saayman et al., 2023), which we found to be specifically in repeats adjected to the alpha satellites (Marselli et al., bioRxiv 2026). When errors accumulate, cells can undergo defects in kinetochore binding and errors in chromosome segregation, leading to numerical and structural alterations that are a major contributor to genetic diseases and various types of cancers (Santaguida and Musacchio, 2009; Gordon et al., 2012; Santaguida and Amon, 2015; Dumont et al., 2020). During replication stress, upon perturbation of centromeric chromatin, such instability at centromere and ensuing DNA damage can lead to mitotic DNA synthesis (MiDAS) and whole arm events such as isochromosomes and chromosomal translocations (Giunta et al., 2021).

Given the physiological dynamics at centromeres, we set out to investigate the impact of exogenous replication stress on these fragile loci. Here, we uncovered a surprising consequence of exposing non-cancerous diploid cell lines from a variety of somatic tissues to replication stress induced by prolonged exposure to low dose hydroxyurea (HU). We found presence of many early G1 doublets, still visibly connected at cytoplasmic level, with one high and one very low amount of γH2AX, suggesting Non-Random Segregation (NRS) of the damage. In previous studies, the NRS phenotype was associated only with differential segregation of the parental DNA strands, the immortal strand hypothesis (Cairns, 1975), where it was observed that cells, in response to replication stress, selectively segregate the chromatids carrying the oldest or newest DNA template (Yadlapalli and Yamashita, 2013; Akera et al., 2017; Ginda et al., 2017; Xing et al., 2020). Due to replication stress, the DNA can accumulate DSBs, with a subsequent phosphorylation of the H2A histone variant H2AX and the accumulation of visible γH2AX foci (Valdiglesias et al., 2013). The accumulation of γH2AX occurs within seconds, spreads over tens of kilobases of DNA flanking the break site, and sustain the immediate initiation of the DNA damage responses (DDR) signaling cascade (Celeste et al., 2003). Cells lacking H2AX present impaired DNA damage repair and display gross chromosomal rearrangements (Celeste et al., 2002). We found that, immediately after mitotic division and prior to full separation of the cytoplasm by cytokinesis, the two daughter nuclei exhibit a significant difference in the amount of γH2AX foci, with one daughter nucleus showing visible signal and the other being virtually damage-free. Using a precise series of cell synchronization protocols optimized for each cell line, we induced replication stress in S phase, accumulated cells in mitosis and then captured them at early G1, before full cytoplasmic separation (Methods). We tested several non-cancerous cell lines, including hTERT RPE-1, hTERT BJ, hCEC D29, hTERT Human Fibroblasts, as well as HCT116 and U2OS cancer cells. Following the induction of replication stress with HU, we observed a NRS of the DNA damage marker γH2AX in all non-cancer cell lines. Importantly, none of the cancer cell lines showed the stress-response phenotype. Data obtained using hTERT RPE-1 cells expressing a Doxycycline-inducible RNaseH1 to remove R-loops, and an Auxin inducible degron to control the degradation of CENP-A, showed that the γH2AX NRS phenotype was increased upon depletion of CENP-A and reduced upon removal of R-loops. This implies that centromere stability is important for the correct orientation and segregation of γH2AX between the two daughter cells. Notably, chemical inhibition of the catalytic activity of Rad51 reduced the NRS of DNA damage, while the total number of damaged cells remains unchanged. These results suggest an involvement of the strand invasion activity of Rad51 in accumulating γH2AX in only one chromatid ahead of cell division. We found that the accumulation of R-loops and DNA damage at the centromere affects the binding of the kinetochore proteins, with measurable effects on the spindle length, orientation, and asymmetry at the metaphase plate. Altogether, we present a new mechanism for preserving genetic information in response to replication stress. We show the unprecedented results that replication stress and/or CENP-A depletion triggers the accumulation of R-loops and DSBs. Selective strand invasion by Rad51 then accumulate γH2AX-labelled foci in one chromatid, leading to spindle asymmetry, whereby the differentially oriented, damaged chromatids are segregated into one daughter cell compared to the undamaged one.

## RESULTS

### Replication stress induces Non-Random Segregation of DNA damage marker γH2AX to one daughter cell

Replication stress is widely recognized as a leading cause of genome instability, with major consequences for cell survival, arising from a variety of sources, such as limiting nucleotides, DNA lesions, repetitive DNA elements, transcription complexes, and/or RNA-DNA hybrids (Zeman and Cimprich, 2014). Any obstacle that perturbs the progression of the replication forks, including affecting components of the replication machinery, is generally considered a form of replication stress that can generate DSBs (Mailand et al., 2013). Moreover, the accumulation of DNA damage resulting from replication stress has been linked to a differential segregation of the parental DNA strands between the daughter cells, also known as the immortal strand hypothesis (Cairns, 1975; Xing et al., 2020).

To investigate the consequences of replication stress and accumulation of R-loops after the mitotic segregation, we set up a mitotic shake-off protocol to induce replication stress and DNA damage specifically during the S phase (Fig. 1A, left). We used thymidine to accumulate hTERT RPE-1 and hTERT BJ cells at the G1/S boundary, followed by release into medium containing HU to induce replication stress and DNA damage accumulation during the S phase, and RO-3306 to generate a second cell cycle block at the G2/M boundary. Finally, cells were released into M phase before performing a mitotic shake-off. The mitotic cells were then collected and re-plated to allow them to reach the G1 phase prior to immunofluorescence analysis (Fig. 1A, right). After HU-induced replication stress, we observed the NRS of the DNA damage marker γH2AX (Fig. 1B). This NRS phenotype was present in ∼40% of the G1 couples across two different batches of RPE-1 cells, and in ∼35% of G1 couples of BJ cells (Fig. 1C). To investigate the overall damage induced after replication stress; we then analyzed the remaining G1 couples. We observed G1 couples in which neither nucleus contained γH2AX-labelled DNA, and G1 couples in which both nuclei presented γH2AX foci (Methods and Fig, 1D). In untreated cells, around 90% of G1 couples were undamaged in both RPE-1 and BJ cells (Fig. 1E), with the remaining ∼10% distributed across the other two categories. Upon perturbation with HU, the percentage of non-damaged couples dropped to ∼20% in the RPE-1 and ∼25% in BJ cells (Fig. 1E).

**Figure 1:**
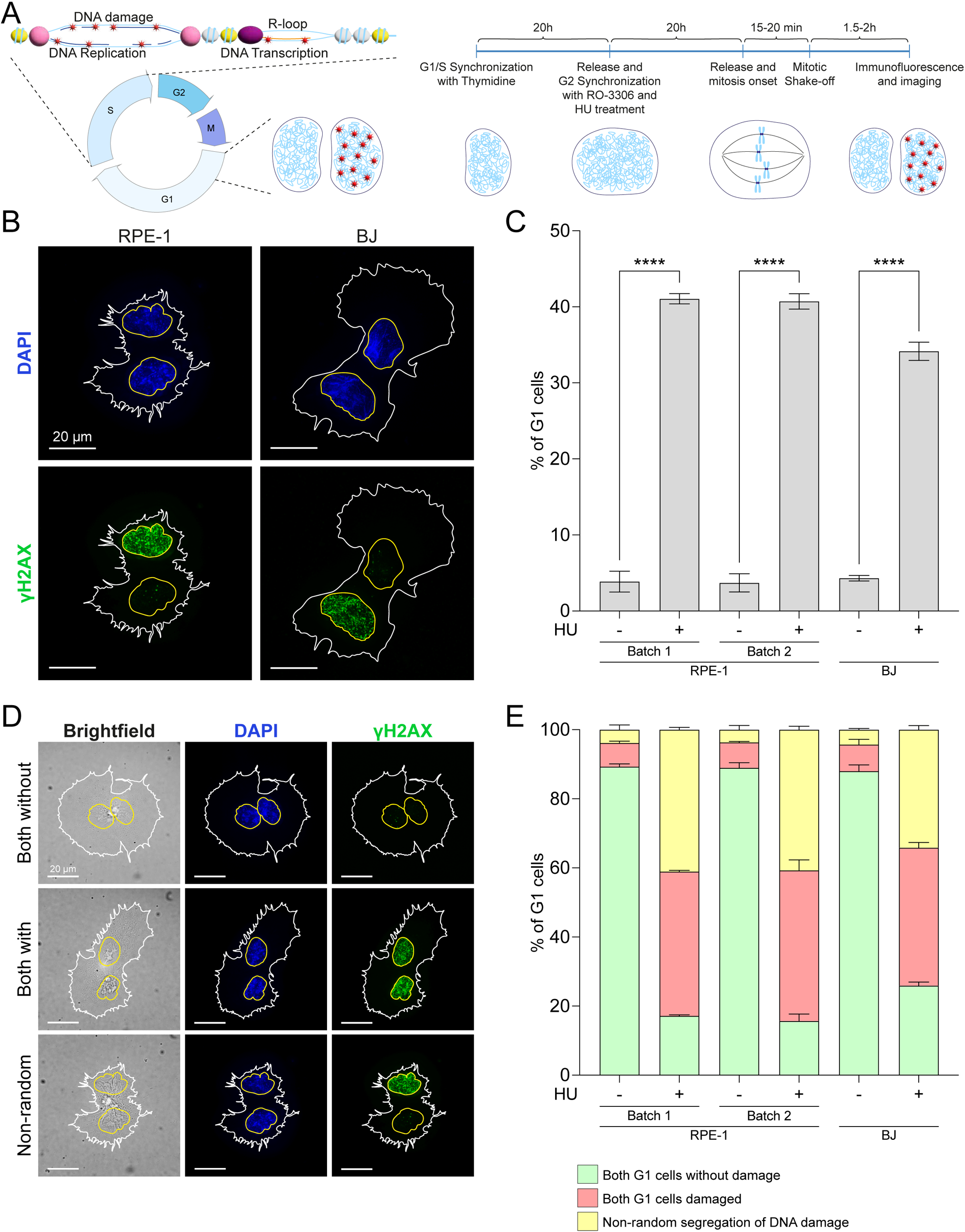
Replication stress induces the Non-Random Segregation of γH2AX. (A) Diagram of the mitotic shake-off protocol used to induce replication stress and DNA damage accumulation specifically during S-phase and to analyze its consequences on mitotic segregation. (B) Examples of G1 couples from RPE-1 (left) and BJ (right) cells showing NRS of DNA damage. The white outline highlights the cytoplasm shape, while the yellow one highlights the nucleus shape, scalebar is 20 µm. (C) Quantification of G1 cells presenting NRS of γH2AX in RPE-1 and BJ cells, with and without HU treatment. For each individual experiment, 55-80 G1 couples were counted. The error bars show the SD of the mean across three biological replicates. *p*-value is from ANOVA test. (D) Examples of RPE-1 cells showing: both G1 without DNA damage, both G1 with DNA damage, NRS of DNA damage. The white outline highlights the cytoplasm shape, while the yellow one highlights the nucleus shape, scalebar is 20 µm. (E) Quantification of the RPE-1 and BJ G1 cells as the examples in D, with and without HU treatment. For each individual experiment, 55-80 G1 couples were counted. The error bars show the SD of the mean across three biological replicates.

To test whether the phenotype can be observed independently to the two consequently synchronizations, we performed two additional mitotic shake-off experiments using different synchronization protocols. In the first setup, we treated RPE-1 cells with HU without any prior synchronization (Asynchronous). In the second, we synchronized RPE-1 cells in G1 with Palbociclib and then added HU upon release to induce replication stress. Similarly to the cells synchronized with Thymidine and RO-3306, we observed the NRS of γH2AX, though at different levels: ∼23% and ∼30% respectively (Fig. 2A). These lower NRS levels are likely due to the differences inherent to the synchronizations protocols themselves, a different number of cells will be damaged by HU depending on whether it is added to asynchronous cells or to synchronized cells at the beginning of the S phase, and a different number of cells will be able to segregate γH2AX non-randomly when comparing asynchronous and G2-synchronized cells.

**Figure 2:**
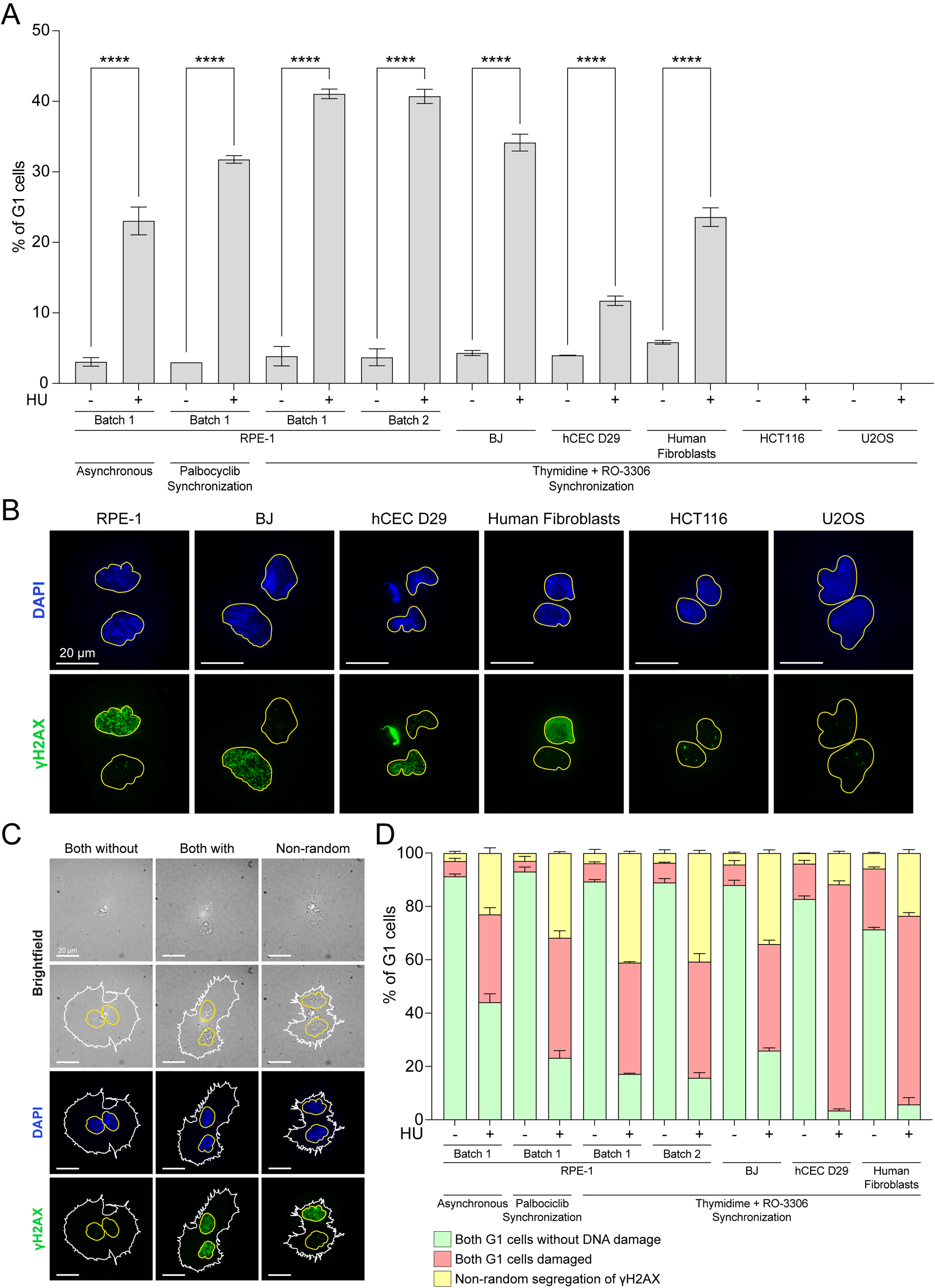
Cancer cells do not manifest Non-Random Segregation of DNA damage. (A) Quantification of G1 cells presenting NRS of γH2AX in RPE-1, BJ, hCEC D29, human fibroblasts, HCT116 and U2OS cells, with and without HU treatment. For each individual experiment, 55-80 G1 couples were counted. The error bars show the SD of the mean across three biological replicates. *p*-value is from ANOVA test. (B) Examples of G1 couples from RPE-1, BJ, hCEC D29 and human fibroblasts cells showing NRS of DNA damage, and HCT116 and U2OS not showing it. (C) Examples of RPE-1 cells showing: both G1 without DNA damage, both G1 with DNA damage, NRS of DNA damage. The white outline highlights the cytoplasm shape, while the yellow one highlights the nucleus shape, scalebar is 20 µm. (D) Quantification of the RPE-1, BJ, hCEC D29 and human fibroblasts G1 cells as the examples in C, with and without HU treatment. For each individual experiment, 55-80 G1 doublets were counted. The error bars show the SD of the mean across three biological replicates.

Next, we investigated the occurrence of NRS in different normal and cancer cell lines (Supplementary Fig. 1 and 2), to determine whether the phenotype was exclusive to certain cell types. For this purpose, human Colon Epithelial Cells (hCEC D29) (Bosco et al., 2023; Mays et al., 2023), immortalized human fibroblasts (hTERT Human Fibroblasts) (Rinaldo et al., 2012), human colorectal cancer cells (HCT116) and human osteosarcoma cells (U2OS) were used alongside RPE-1 and BJ cells. Overall, we observed that high levels of DNA damage and replication stress, induced by HU treatment across all cell lines, triggered the NRS of γH2AX labelled DNA in all normal but not in cancer cell lines (Fig. 2A and Fig. 2B). Among the non-cancer cell lines, we observed variability in the occurrence of NRS, going from ∼15% in hCEC D29, to ∼40% in RPE-1 (Fig. 2A and 2B). We also observed variability in the overall accumulation of DNA damage, with different proportions of cells presenting γH2AX in both G1 daughter nuclei (Fig. 2C and 2D, Supplementary Fig. 1 and 2). Together, these data indicate that different cellular backgrounds can influence the response to replication stress and DNA damage, specifically in physiological cells. Overall, in line with previous findings that NRS can trigger a differential DNA damage response between the parental templates leading to differential distribution of damaged DNA in the daughter cells (Xing et al., 2020), our data reinforce this idea by adding the notion that replication stress triggers the NRS of DNA damage markers already under physiological conditions.

### Centromeric R-loops removal affects Non-Random Segregation of DNA damage

One of the main historical consequences of replication stress is the occurrence of breaks and gaps on mitotic chromosomes at the Common Fragile Sites (CFSs) (Glover et al., 1984; Debatisse et al., 2012). CFSs are normal components of the chromosome structure that are prone to instability due to a combination of delayed replication (Le Beau et al., 1998; El Achkar et al., 2005), transcription of long genes (Smith et al., 2006; Bosco et al., 2010) and defective condensin resulting from an under-replicated state that persists until mitosis (Boteva et al., 2020). Replication stress leads to the accumulation of R-loops and DNA damage following the collision of replication and transcription machineries (Helmrich et al., 2011); at the centromere, the accumulation of R-loops has been linked to chromosome breaks and translocations (Giunta et al., 2021). These features, among others, make the centromeres similar to CFSs (Black and Giunta, 2018). Like CFSs, centromeres are also characterized by late replication, peaking at 5 h after satisfaction of the G1/S checkpoint and continuing into G2 (Ten Hagen et al., 1990; Massey et al., 2019); active transcription (McNulty et al., 2017); recombination (Giunta and Funabiki, 2017); and by a propensity to form non-B-DNA and secondary structures (Kabeche et al., 2018; Kasinathan and Henikoff, 2018; Chittoor and Giunta, 2024).

To investigate the relationship between NRS and the effects of R-loops accumulation and removal (Fig. 3A), we used RPE-1 cells expressing Doxycycline (Dox)-inducible RNaseH1 (Giunta et al., 2021). Efficiency of RNaseH1 induction was confirmed by immunofluorescence detection of co-expressed green fluorescent protein (GFP) (Supplementary Fig. 3). Specifically, two different cell line batches were used: one expressing a form of RNaseH1 capable of targeting the whole genome (Genomic RNaseH1), and a second expressing an RNaseH1 fused to the CENP-B DNA binding domain (Centromeric RNaseH1). This distinction allowed us to selectively target centromeres and thereby infers the role of R-loops at centromeres in NRS (Supplementary Fig. 3). We used the same mitotic shake-off protocol as before (Fig. 1A): upon releasing the cells from thymidine, we added HU to induce replication stress, together with Dox to promote R-loops resolution. In both RPE-1 cell batches, activation of RNaseH1 to remove HU-induced R-loops led to a slight reduction in the number of G1 couples showing NRS of γH2AX (Fig. 3B). To determine whether the effect of R-loops removal was related to the total amount of DNA damage or to its segregation, we analyzed the remaining G1 couples (Fig. 3C, bottom). Removal of R-loops did not change the total amount of DNA damage but rather affected its segregation between the two daughter cells (Fig. 3D). We further investigated the contribution of R-loops to the observed NRS differences by analyzing the DNA damage in the G2 cells. We applied the same synchronization protocol used for the mitotic shake-off, but we fixed the cells before releasing the RO-3306 block (Fig. 3C, top). As observed in G1 cells, there were no differences in the total DNA damage with or without the removal of R-loops induced by replication stress (Fig. 3D).

**Figure 3:**
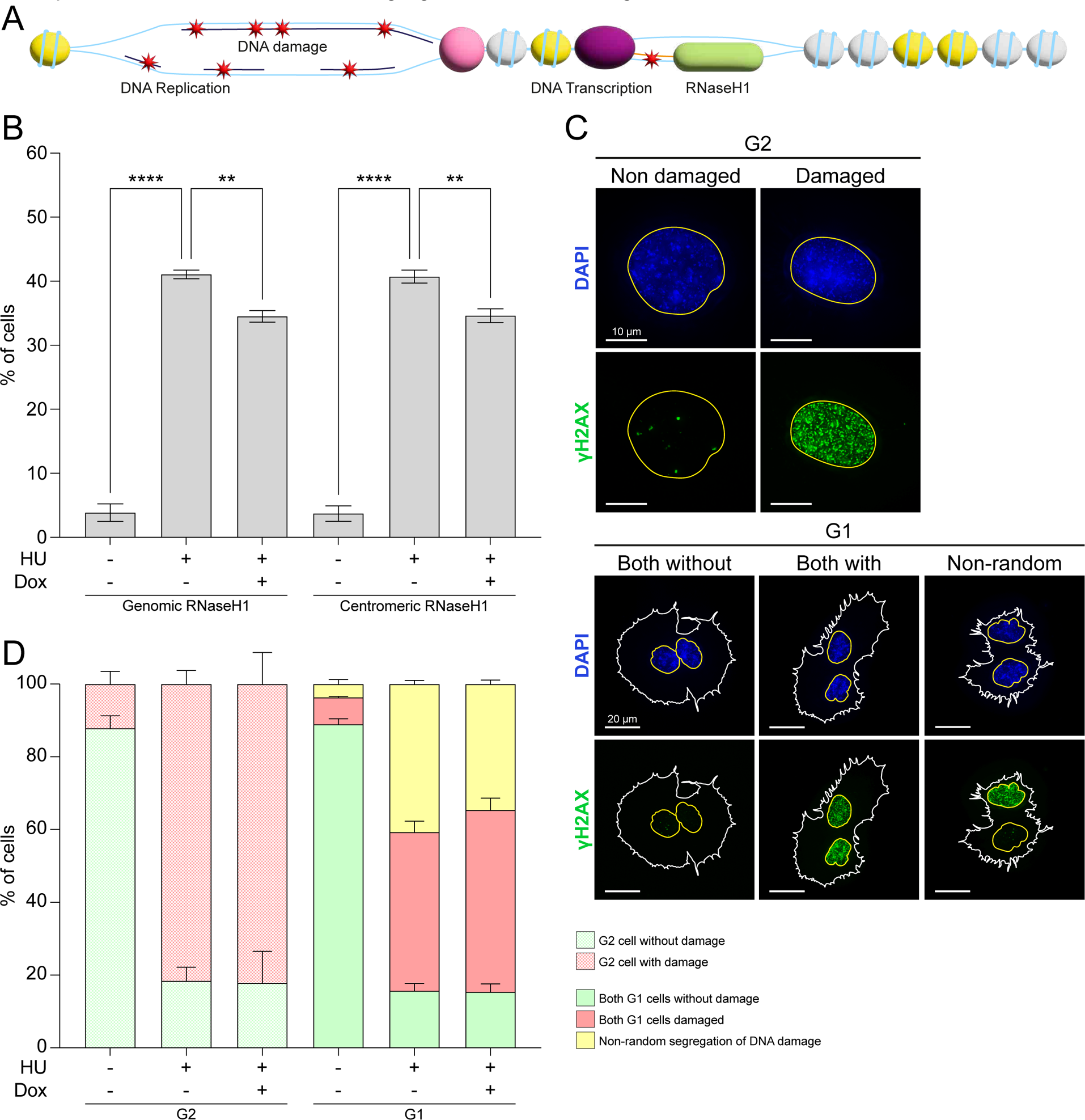
R-loops contribute to Non-Random Segregation of DNA damage. (A) Diagram of centromeric chromatin showing the accumulation of DNA damage after replication stress and RNaseH1 binding to R-loops. (B) Quantification of G1 cells presenting NRS of γH2AX in Genomic and Centromeric RNaseH1 RPE-1 cells, with HU and Dox treatments. For each individual experiment, 55-80 G1 couples were counted. The error bars show the SD of the mean across three biological replicates. (C) On top, examples of G2 cells showing: G2 without DNA damage and G2 with DNA damage, scalebar is 10 µm. On bottom examples of RPE-1 cells showing: both G1 without DNA damage, both G1 with DNA damage, NRS of DNA damage. The white outline highlights the cytoplasm shape, while the yellow one highlights the nucleus shape, scalebar is 20 µm. (D) On the left side of the graph, quantification of the RPE-1 G2 cells as the examples in C, with and without HU and Dox treatments. For each individual experiment, 100-120 G2-arrested cells were counted. On the right side of the graph, quantification of the RPE-1 G1 cells as the examples in C, with and without HU and Dox treatments. For each individual experiment, 55-80 G1 couples were counted. The error bars show the SD of the mean across three biological replicates.

Because the removal of R-loops slightly reduced the NRS following replication stress without affecting overall DNA damage (Fig. 3D), irrespective of whether Centromeric or Genomic RNaseH1 was used, these data indicate that the presence of these DNA-RNA hybrids at the centromere could affect the stability and/or binding of the kinetochore, leading to a non-random orientation of chromosomes on the metaphase plate.

### CENP-A depletion induces Non-Random Segregation of DNA damage

To ensure the correct segregation of sister chromatids during mitosis, it is crucial for the kinetochore to properly bind centromeric chromatin, a process mediated by CENP-A (Schueler and Sullivan, 2006; Santaguida and Musacchio, 2009; Hayden, 2012; Barra and Fachinetti, 2018; Altemose et al., 2022). The accumulation of R-loops at centromere, followed by centromere instability and breaks, has been linked to the destabilization of CENP-A (Giunta et al., 2021).

The RPE-1 cells used in this study also carry an Auxin (AUX)-inducible degron construct targeting CENP-A (Giunta et al., 2021), an important feature for studying the role of CENP-A and centromere stability in the NRS phenotype. Depletion of the centromeric nucleosome was monitored via the YFP fluorophore fused to CENP-A (Supplementary Fig. 4). We induced CENP-A depletion during S phase by adding AUX together with HU and RO-3306 (Fig. 4A, Supplementary Fig. 4), used to induce the accumulation of R-loops and DNA damage specifically at the centromere. We observed NRS in ∼25% of cells upon CENP-A depletion alone, highlighting that centromere stability is crucial for the occurrence of NRS. Interestingly, CENP-A depletion had an additional detrimental effect on top of HU-induced replication stress: indeed, CENP-A depletion increased HU-induced NRS by 10% (50% CENP-A AID + HU vs. 40% HU only) (Fig. 4B). Because the effect of centromere instability did not increase NRS to the expected ∼65% (40% HU + 25% AUX, which would indicate a fully additive effect), this suggests that HU is already partially affecting centromere stability, consistent with centromeres behaving similarly to CFSs (Black and Giunta, 2018). As for replication stress, also CENP-A depletion causes the accumulation of DNA-RNA hybrids (Giunta et al., 2021). Accordingly, to investigate their role in NRS, we combined CENP-A depletion with RNaseH1 expression (Fig. 4A). Similarly to the reduction in NRS observed after R-loops removal in HU damaged cells, we observed a comparable effect upon removing R-loops induced by CENP-A depletion alone, or in combination with HU-inducible replication stress (Fig. 4B).

**Figure 4:**
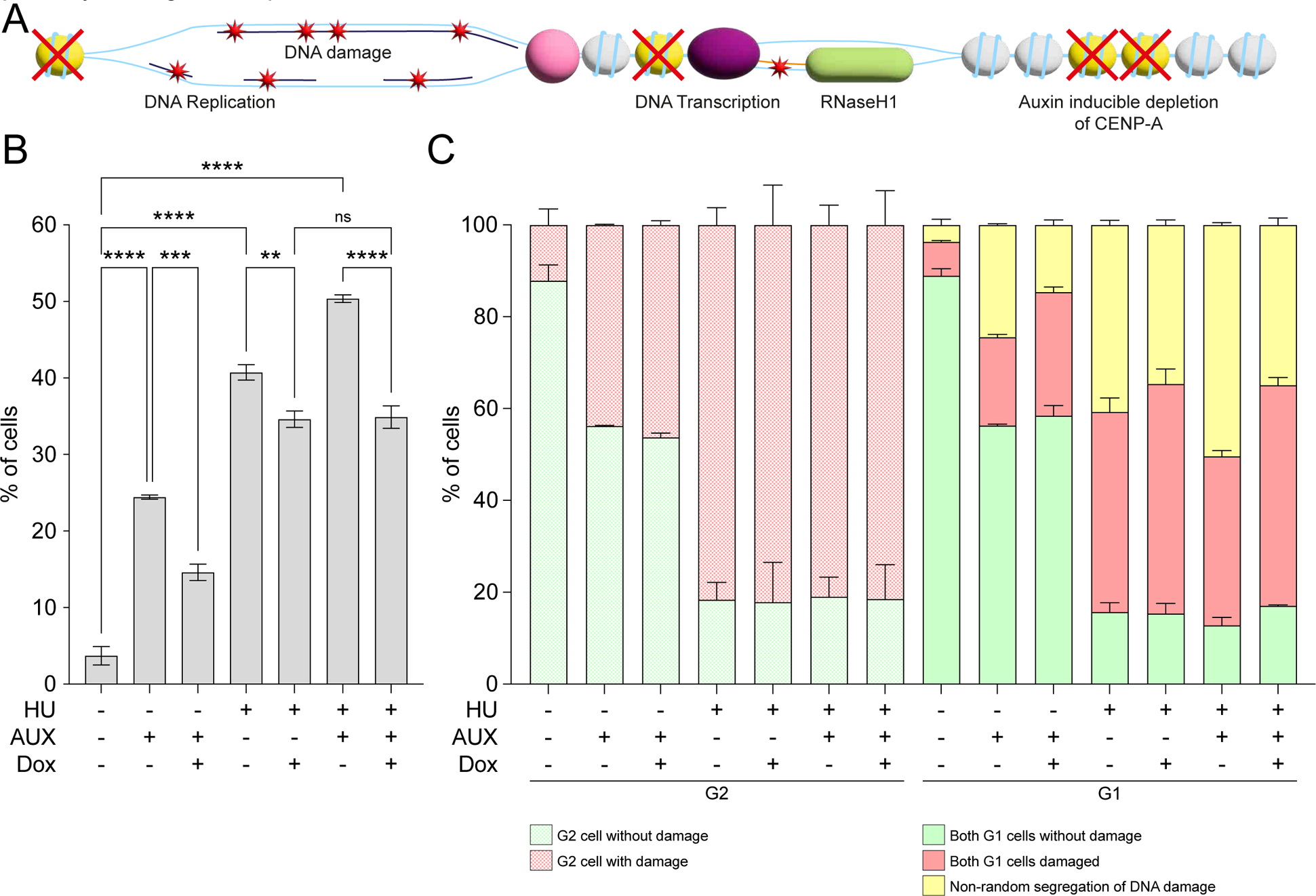
Centromere specific H3 histone variant CENP-A depletion causes Non-Random Segregation of DNA damage. (A) Diagram of centromeric chromatin showing the accumulation of DNA damage after replication stress and CENP-A depletion, and RNaseH1 binding to R-loops. (B) Quantification of G1 cells presenting NRS of γH2AX in Centromeric RNaseH1 RPE-1 cells, with HU, AUX and Dox treatments. For each individual experiment, 55-80 G1 couples were counted. The error bars show the SD of the mean across three biological replicates. (C) On the left side of the graph, quantification of the RPE-1 G2 cells as the examples in 3C, with and without HU, AUX and Dox treatments. For each individual experiment, 100-120 G2s were counted. On the right side of the graph, quantification of the RPE-1 G1 cells as the examples in 3C, with and without HU, AUX and Dox treatments. For each individual experiment, 55-80 G1 couples were counted. The error bars show the SD of the mean across three biological replicates.

When analyzing the total G1 cell population, we did not observe changes in the total amount of DNA damage upon CENP-A depletion, with or without concomitant R-loop removal (Fig. 4C, Supplementary Fig. 5). Moreover, we did not observe changes in the total DNA damage induced by replication stress alone or in combination with CENP-A depletion (Fig. 4C, Supplementary Fig. 5). We further investigated the effect of CENP-A depletion on NRS variation by analyzing cells in G2 prior to mitotic segregation (Fig. 3C) (see previous section). As in G1 cells, we did not observe changes in the total DNA damage when comparing cells after CENP-A depletion, with or without the R-loop removal. Similarly, there were no differences upon HU-induced replication stress, alone or in combination with CENP-A depletion, in the presence or absence of R-loops (Fig. 4C, Supplementary Fig. 5).

The NRS observed as consequence of CENP-A depletion strengthens the notion that the stability or instability of the centromere, and/or the presence or absence of R-loops, create a differential marking of the kinetochore binding site, driving either random or non-random segregation of DNA damage between the two daughter cells.

### Rad51 contributes to asymmetric DNA damage distribution

After replication stress induced by HU, the replication fork stalls due to inhibition of nucleotide incorporation. Prolonged stalling of replication forks can eventually lead to the accumulation of DSBs. The restart of stalled replication forks is a fundamental driving force behind *Escherichia coli* fitness, where reactivation of replication dictates viability due to the absolute lack of backup origins (Heller and Marians, 2006). In mammalian cells, agents that cause stalling or collapse of replication forks, such as HU, strongly induce the activation of the Homologous Recombination (HR) DNA damage repair pathway, which promotes cell survival following these treatments (Arnaudeau et al., 2001; Saintigny et al., 2001; Lundin et al., 2002), allowing restoration of the DNA molecule during S and G2 phases (Li and Heyer, 2008) (Fig. 5A). Accordingly, one of the main DNA repair factor ensuring the smooth progression of DNA replication forks is RAD51, which is critical not only for the HR, but also for the fork reversal pathway (Quinet et al., 2017; Bhat and Cortez, 2018; Bhowmick et al., 2022). Mechanistically, Rad51 binds single stranded DNA (ssDNA) to form the RAD51-ssDNA nucleofilament (Esashi et al., 2007) and search for the homologous DNA sequences to invade and use as templates.

**Figure 5:**
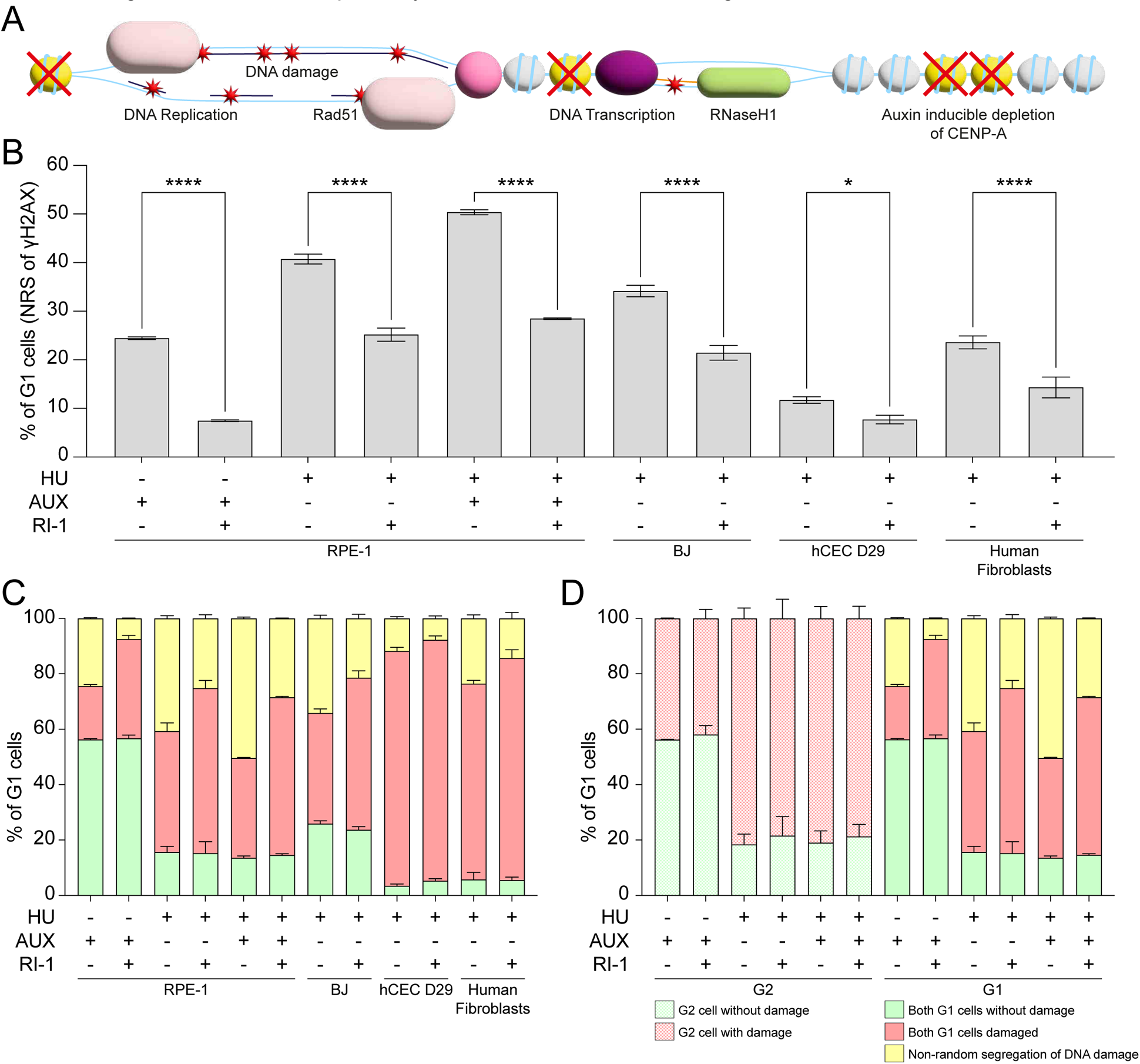
The homologous recombination pathway accumulates the replicative DNA damage on one chromatid. (A) Diagram of centromeric chromatin showing the accumulation of DNA damage after replication stress and CENP-A depletion, RNaseH1 binding to R-loops and Rad51 binding to DNA damage. (B) Quantification of G1 cells presenting NRS of γH2AX in RPE-1, BJ, hCEC D29 and Human Fibroblasts with and without HU, AUX and RI-1 treatments. For each individual experiment, 55-80 G1 couples were counted. The error bars show the SD of the mean across three biological replicates. *p*-value is from ANOVA test. (C) Quantification of the RPE-1, BJ, hCEC D29 and human fibroblasts G1 cells as the examples in 1C, with and without HU, AUX and RI-1 treatments. For each individual experiment, 55-80 G1 couples were counted. The error bars show the SD of the mean across three biological replicates. (D) On the left side of the graph, quantification of the RPE-1 G2 cells as the examples in 3C, with and without HU, AUX and RI-1 treatments. For each individual experiment, 100-120 G2s were counted. On the right side of the graph, quantification of the RPE-1 G1 cells as the examples in 3C, with and without HU, AUX and RI-1 treatments. For each individual experiment, 55-80 G1 couples were counted. The error bars show the SD of the mean across three biological replicates.

To assess the involvement of Rad51 in the NRS phenotype, and, more specifically to determine whether the strand-invasion activity of Rad51 could be important in the NRS, potentially by driving accumulation of γH2AX in only one of the two sister chromatids, we performed loss-of-function studies by inhibiting Rad51 with RI-1. The chemical inhibitor RI-1 binds covalently to cysteine 319 on the surface of the Rad51 protein, inactivating it by blocking the interface used for protein-protein interactions between subunits (Budke et al., 2012). We applied the inhibitor to all the normal cell lines under all DNA damage conditions examined in this study, CENP-A depletion, replicative stress, or both combined, and in each case we observed a strong reduction in the proportion of G1 couples showing NRS of γH2AX (Fig. 5B). As before, we analyzed all the G1 couples to capture the full range of possible γH2AX segregation patterns, which showed that the reduction in NRS was not due to a reduction in total DNA damage, but rather to a different pattern of accumulation in the two sister chromatids: in all cell lines, we observed an increased number of couples in which both G1 nuclei were damaged (Fig. 5C). As for the NRS reduction observed after R-loops removal, we verified this by analyzing the cells in G2 phase (Fig. 5D and Supplementary Fig. 6), confirming that the difference in NRS is not due to differences in the total DNA damage.

While RI-1 is generally used at a concentration between 1 and 50 μM to inhibit HR, we selected a concentration of 20 μM. To investigate the potential dose-dependency of the NRS reduction in G1 cells, further experiments were conducted at the outer limits of this range, including then the lowest (1 μM) and highest (50 μM) concentrations. We observed a greater reduction in NRS with increasing doses of RI-1, up to 50 μM. As before, there was no difference in the total amount of DNA damage between cells under replication stress alone, and cells co-treated with the Rad51 inhibitor (Supplementary Fig. 7A and 7B).

Taken together, our data show that the HR protein Rad51 is activated in response to DNA damage accumulation induced by HU and/or CENP-A depletion. Because we observed no difference in the total DNA damage with or without Rad51 activity, we propose that Rad51 acts independently of the DNA damage repair pathway. It has been previously demonstrated that Rad51 activity is pivotal for resolving the stalled replication forks induced by HU without activation of HR (Petermann et al., 2010). This alternative exchange pathway is likely responsible for the redistribution of γH2AX in one chromatid, leaving the other undamaged, so that the cell can segregate the damage non-randomly.

### Spindle asymmetry reveals influence on kinetochore binding and ensuing division

To determine what could be the discriminating factor that distinguishes damaged from non-damaged chromatids in driving NRS. Centromeric DNA is associated throughout the cell cycle with the Constitutive Centromere-Associated Network (CCAN), a macromolecular complex that bridges CENP-A with the outer layer of the kinetochore through the activity of CENP-C (Di Tommaso and Giunta, 2023). Mitotic centromere transcription is important for stabilizing the interaction of CENP-C with the kinetochore, although transcription of alpha-satellites, occurring concomitantly with their replication, is limited by a still-unknown CENP-A-dependent mechanism (Chan and Wong, 2012; Chan et al., 2012). As a consequence of CENP-A depletion, as well as of replication stress, collision between the transcription and replication machineries generate mutagenic R-loops that are deleterious to centromere stability (Giunta et al., 2021). The generation of mutagenic R-loops and DNA damage at centromeres sets in motion a chain reaction of disruptive events that undermine centromere function and the checkpoint ensuring proper microtubule attachment for correct chromosome segregation (Stern and Murray, 2001; Liu and Lampson, 2009; Musacchio, 2015; Papini et al., 2021), potentially impairing the binding of kinetochore proteins.

Based on this, we hypothesized that one of the mechanisms dictating NRS of γH2AX might be a differential binding of kinetochore proteins between the sister chromatids, as a consequence of the accumulation of DNA damage and R-loops. This altered binding would result in differences in microtubule pulling forces between damaged and non-damaged chromatids, leading to an orientation of chromosomes on the metaphase plate that drives NRS (Fig. 6A). To analyze the behavior of the mitotic spindle, we synchronized RPE-1 cells at the G2/M boundary with RO-3306 for 24 hours, then released them to accumulate mitotic cells, which were fixed with paraformaldehyde prior to immunofluorescence staining. For each mitotic cell, we measured the length of the mitotic spindles on both sides, from the pole to the centromeres, and scored the metaphases as damaged or non-damaged based on γH2AX staining (Fig. 6B). Under replication stress, we observed an overall increase in the mean spindle length (from ∼4 to ∼5 μm) compared to untreated cells (Fig. 6C). We then calculated the absolute value of the difference between the two spindle sides (Methods): a low value indicates that the two sides are similar in length, while a high value indicates that they differ. In cells that experienced replication stress, we observed higher spindle length differences compared to untreated cells, particularly in metaphases displaying DNA damage (Fig. 6D), also reflected in the slope of the line connecting the measurements of the two sides (Supplementary Fig. 8). These data show that the accumulation of DNA damage and R-loops lead to the instability of kinetochore binding, affecting the microtubule pulling forces.

**Figure 6:**
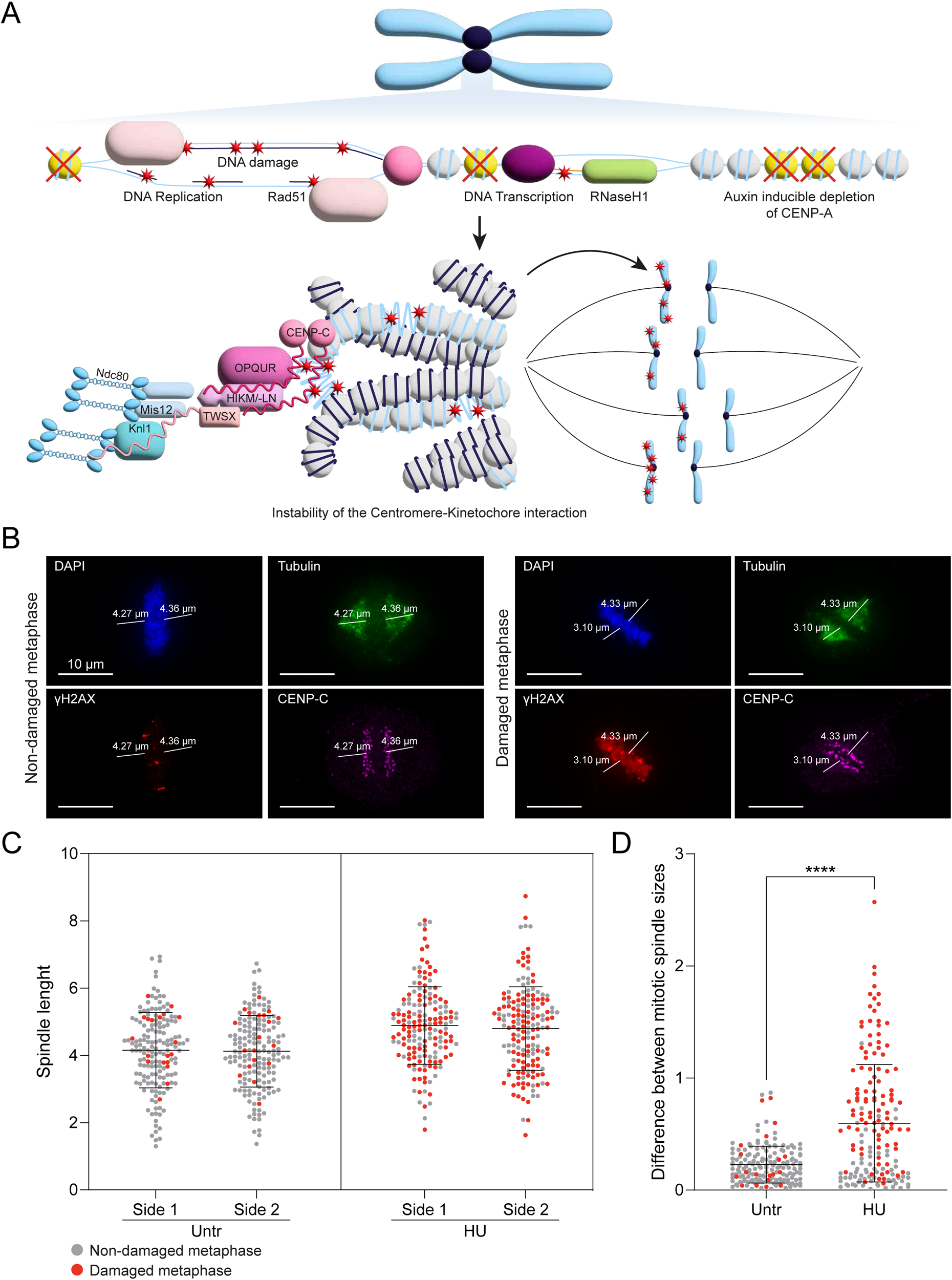
Kinetochore binding and epigenetic marks promote differential segregation of damaged chromatids. (A) Diagram of centromeric chromatin showing the accumulation of DNA damage after replication stress and CENP-A depletion, RNaseH1 binding to R-loops and Rad51 binding to DNA damage. The presence of mutagenic R-loops, DNA damage, and the absence of CENP-A can affect the binding of the kinetochore proteins differentiating the damaged and the non-damaged chromatids with a consequent difference in the microtubule pulling forces. (B) Examples of non-damaged and damaged metaphases with tubulin staining to measure the size of the mitotic spindle. (C) Quantification of both sides of the spindle in unperturbed cells and mitosis after replicative stress, color scheme as in B. For each condition, ∼130 metaphases were counted. The error bars show the SD of the mean. (D) Quantification of the difference between the two sides of the spindle in unperturbed cells and mitosis after replicative stress, grey dots are non-damaged metaphases, red dots are damaged metaphases. For each condition, ∼130 metaphases were counted. The error bars show the SD of the mean.

Overall, we describe here how, following replication stress, there is an alteration in kinetochore protein binding. The presence of γH2AX on one sister chromatid, mutagenic R-loops and the instability of CENP-A, all contributes to the destabilization of the interaction with CENP-C (Di Tommaso and Giunta, 2023). The differential kinetochore interaction occurring in damaged versus non-damaged chromatids is then translated into differences in microtubule pulling forces, leading to a non-random orientation of chromatids on the metaphase plate (Fig. 6A).

## DISCUSSION

Here, we observed the Non-Random Segregation of the DNA damage marker γH2AX after exposing cells to replication stress. The bias toward segregating damage into just one of the two daughter cells upon mitotic exit was observed exclusively in non-cancer human cell lines, and it does not occur under the same perturbations in any of the cancer cell line analyzed here. We used HU to induce DNA damage and R-loops formation in fragile regions of the genome (Glover et al., 1984; Debatisse et al., 2012), including centromeres (Black and Giunta, 2018; Giunta et al., 2021). We applied a series of synchronizations to selectively trigger instability during S phase and then study the cell’s response upon mitotic exit, in the following G1 phase (Fig. 1A). During replication stress, CFSs (Maccaroni et al., 2020; Balzano et al., 2021) and centromeres (Black and Giunta, 2018; Balzano et al., 2021) are among the most affected regions, as they are prone to accumulate DNA damage and to delay replication into G2 phase (Glover et al., 1984; Debatisse et al., 2012; Massey et al., 2019; Scelfo et al., 2024). After replication stress, the non-cancer cell lines used in this study, RPE-1, BJ, hCEC D29, and human fibroblasts, showed a significant difference in the amount of γH2AX observed between the two daughter cells (Fig. 1B, Fig. 2B, Supplementary Fig. 1, Supplementary Fig. 2). This is in line with the NRS phenotype observed for the Cyclobutane Pyrimidine Dimers (CPDs) following UV irradiation at the G2/M boundary, potentially reflecting a local asymmetry at the histone level between sister chromatids (Ferrand et al., 2025). In addition, our results reveal a broader chromosome-specific mechanical layer that relies on the segregation machinery after the most common damage, the replication stress. Here, we found that this phenotype correlates with an accumulation of R-loops at centromeres following replication stress, creating an asymmetry in kinetochore binding and activity that can bias segregation. We further investigated this relationship between centromere instability and biased damage segregation by inducing CENP-A depletion (Supplementary Fig. 4), which we have previously shown to lead to accumulation of detectable R-loops (Giunta et al., 2021). Using two inducible RNaseH1 systems, one global and one centromere-specific (Supplementary Fig. 3), we were able to resolve this R-loop accumulation. Under these conditions, we found that centromeric instability induced by CENP-A depletion is sufficient to bias damage transmission and trigger the NRS, and that removal of R-loops partly reduces the NRS (Fig. 3, Fig. 4, Supplementary Fig. 5). These results indicate that the presence of DNA-RNA hybrids and/or the absence of CENP-A promotes NRS of DNA damage, potentially by impairing the binding of kinetochore proteins. It is tempting to speculate that this mechanism could represent a response aimed to preserve genome integrity, by accumulating the damage in only one of the daughter cells while preserving the genetic stability of the other one, as previously proposed in the immortal strand hypothesis (Cairns, 1975; Charville and Rando, 2013; Xing et al., 2020) and shown in stem cells (Potten et al., 2002; Karpowicz et al., 2005; Smith, 2005; Shinin et al., 2006; Armakolas and Klar, 2007; Conboy et al., 2007).

Additionally, we observed that the strand invasion activity of Rad51, important in both HR and replication fork restart pathways, contributes to the establishment of NRS in response to HU-induced replication stress. We tested this hypothesis by inhibiting the catalytic activity of Rad51; impeding the strand invasion led to a reduction in the proportion of G1 cells showing NRS of γH2AX (Fig. 5 and Supplementary Fig. 6). Because this damage remained unrepaired, this suggests that Rad51 is activated independently of HR to resolve the stalled replication forks (Petermann et al., 2010), highlighting an important role for Rad51 in accumulating DNA damage in one chromatid, enabling its non-random segregation. To better dissect the role played by Rad51, it will be important to test Rad51 mutants carrying alterations in different functional domains. This will help clarify which activity, and consequently which pathway, is required to distribute γH2AX prior to mitotic segregation, and will further elucidate the underlying mechanism. Finally, we verified that the impaired CENP-C binding, resulting from R-loop accumulation and reduced centromeric RNA, together with auxin-induced CENP-A depletion, leads to a differential kinetochore binding in the presence or absence of DNA damage, altering the centromere structure we have previously described (Di Tommaso and Giunta, 2023; Di Tommaso et al., 2023). We analyzed the mitotic spindle length in damaged mitosis and observed an increasing asymmetry between the two sides of each metaphase plate, reflecting a difference in kinetochore protein binding in presence or absence of damage (Fig. 6). The details of this impaired binding remain to be elucidated, in particular whether the damaged chromatids are segregated towards the longer or the shorter side of the spindle. Moreover, another important aspect to investigate is the dynamicity of the process, from the onset of the replication stress to the accumulation of the DNA damage in one chromatid, through mitotic segregation, and into the subsequent cell cycle, in order to determine the fate of the affected cell. To this end, a crucial next step will be a time-course experiment using live-cell imaging to dissect all the steps leading to the NRS of γH2AX.

We propose a mechanism activated in response to replication stress and the accumulation of DNA damage during the S and G2 phases. Although the replication machinery is highly accurate, the fidelity of the process is often threatened by exogenous or endogenous stresses, resulting in altered replication fork progression, reduced replication fidelity, and DNA breaks (Zeman and Cimprich, 2014). In response to the accumulation of DNA damage induced by replication stress, the cell activates a safeguarding response that may serve to preserve genome integrity (Fig. 7). The accumulation of stalled replication forks can eventually lead to the their collapse into DSBs (Saintigny et al., 2001; Petermann et al., 2010), accompanied by elevated γH2AX signaling (Celeste et al., 2003). To safeguard genome stability, we propose that the cell activates a mechanism relying on Rad51 and its strand invasion activity. As previously observed (Petermann et al., 2010), Rad51 can be activated to resolve stalled replication forks independently of HR activation; the resulting movement of DNA to restart the stalled forks leads to a corresponding movement of DNA damage into only one of the two sister chromatids (Fig. 7). As cells approach mitosis, the activity of the two sister centromeres becomes destabilized, as the presence or absence of DNA damage, R-loops, and CENP-A all contribute to destabilizing the binding of CENP-C (Chan and Wong, 2012; Chan et al., 2012; Di Tommaso and Giunta, 2023), and consequently of the kinetochore proteins. This instability creates an imbalance between the damaged and the non-damaged chromatids (Fig. 7). This asymmetry in kinetochore binding is then transmitted to the microtubules and to their pulling forces; the resulting asymmetric pulling of chromatids leads to an orientation of chromosomes on the metaphase plate that ultimately result in NRS of DNA damage (Fig. 7).

**Figure 7:**
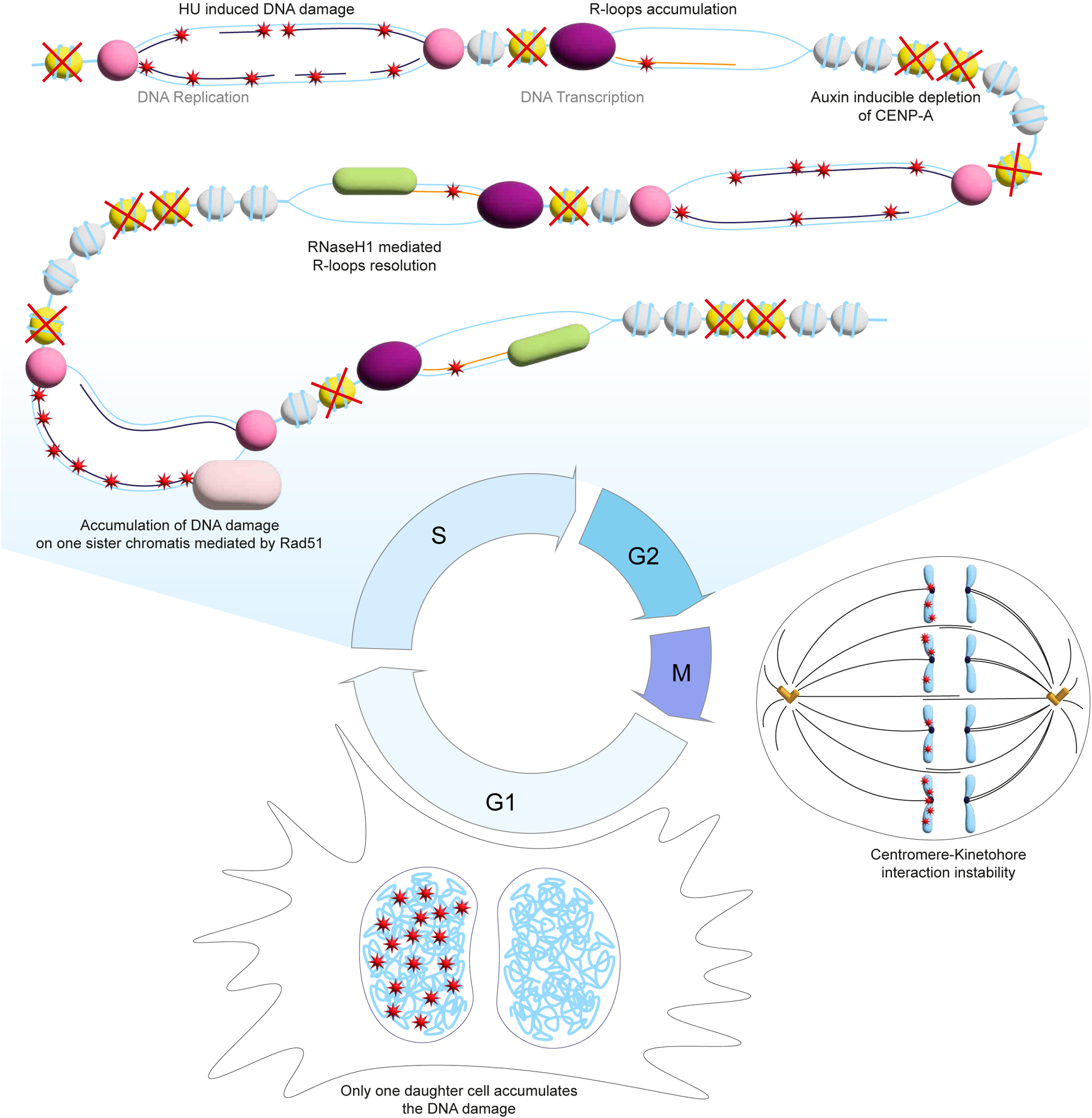
Mechanistic insights into Non-Random Segregation of replication and R-loops induced DNA damage. Diagram showing the proposed mechanism that leads to NRS of DNA damage. HU causes the accumulation of DNA damage and R-loops via replication stress, at centromere level, this is enhanced by the depletion of CENP-A and is mitigated by the activity of the RNaseH1. Rad51 is activated in response to HU, the resolution of the stalling forks has as a consequence the accumulation of the DNA damage to one sister chromatid. Finally, the DNA damage, R-loops, and presence or absence of CENP-A signal the differentiation between the damaged and non-damaged chromatid by interfering with the binding of kinetochore proteins. All of this result in the accumulation of DNA damage in only one of the two daughter cells.

Differential segregation between the older and newer templates has been shown (Cairns, 1975; Xing et al., 2020). Charville and Rando (2013) proposed the “mortal strand hypothesis” according to which, under conditions of replication stress, the older DNA template (immortal strand) and the newer DNA template (mortal strand) accumulate DNA differently, such that sister chromatids may exhibit molecular features reflecting the relative age of their strand. This differential marking of the two chromatids is thought to communicate with the mitotic spindle, promoting or inhibiting the attachment of spindle microtubules to a given chromatid (Charville and Rando, 2013). Asymmetric histone variants distribution(Petryk et al., 2018; Wenger et al., 2023), leading and lagging strands differential accumulation of damage under stress (Charville and Rando, 2013), recombination events and non-random segregation would inevitably be reflected in the daughter cells. Altogether, our work provides new evidence linking the replication stress response to NRS of DNA damage, highlighting a fundamental role for Rad51 pathway in the γH2AX accumulation, as well as a role for the centromere in its mitotic orientation.

## MATERIALS AND METHODS

### Cell culture

All the cell lines used for this project were maintained at 37 °C in a 5% CO2 atmosphere. hTERT RPE-1 modified with an amino-terminal enhanced yellow fluorescent protein (EYFP) and an auxin inducible degron (AID) tags to endogenous CENP-A to facilitate its rapid and complete depletion, and with a Doxycycline inducible RNaseH1 construct fused to a green fluorescent protein, alone or with the CENP-B DNA binding domain to target the centromeres (Giunta et al., 2021). These cells were cultured in DMEM-F-12 supplemented with 10% FBS Tet system approved (GIBCO, A4736201), 1X antibiotic-antimycotic solution (Corning, 30-004-CI), and 2 mM L-glutamine (EuroClone, ECB3000D). Human hTERT immortalized fibroblast cells (hTERT BJ), human colon epithelial cells (hCEC D29), hTERT immortalized human fibroblasts (hTERT human fibroblasts), and colorectal cancer cells (HCT116) were cultured in DMEM High Glucose w Sodium Pyruvate (EuroClone, ECB7501L) with 10% FSB (GIBCO, A5256701), 1X antibiotic-antimycotic solution (Corning, 30-004-CI), and 2 mM L-glutamine (EuroClone, ECB3000D). U2OS cells were cultured in DMEM High Glucose w Sodium Pyruvate (EuroClone, ECB7501L) with 10% FSB (Tet system approved (GIBCO, A4736201), 1X P/S (EuroClone, ECB3001D), and 2 mM L-glutamine (EuroClone, ECB3000D).

### Cell synchronization for mitotic shake-off

Cells are plated in 10 cm dishes at a proper concentration to let them grow for 2 days. For the mitotic shake-off experiments different combinations of synchronizations have been applied. For the main experiments, 40 hours before the harvesting, cells are treated with 2 mM Thymidine to block the cells before entering the S-phase. After 20 hours, the medium with the thymidine is removed and cells are treated with fresh medium supplemented with 9 μM RO-3306 to block the cell cycle at the G2/M border. At this stage, any additional drug of interest is supplemented in the culture medium. After additional 20 hours, cells are released from the G2/M block with fresh medium. After 20 to 40 minutes (depending on the cell line), cells are in mitosis. At this stage, the dishes can be shaken allowing the mitotic cells to detach. The medium containing the mitotic cells is then collected and plated on poly-lysine coverslips. Cells can now divide and enter the G1 phase; the optimal time to accumulate G1 cells depends on the cell line. When there are enough G1 cells, the coverslips can be fixed with 2% paraformaldehyde. One control experiment was then performed with asynchronous cells; the hydroxyurea was added 24h prior the shake-off. A second control experiment was performed synchronizing the cells in G1 with 150 nM Palbociclib, after 20 hours, the medium is removed and cells are treated with hydroxyurea 24h before the mitotic shake-off.

### Cell synchronization of analysis of G2 cells

Cells were plated in 5 cm dishes with 24×24 mm square poly-lysine coverslips at a proper concentration to let them grow for 2 days and treated with Thymidine and RO-3306 as for the mitotic shake-off, but the coverslips were removed from the dish to fix the cells without releasing them from the G2 block.

### Cell synchronization for mitotic cells accumulation

For the analysis of the length of the mitotic spindle cells are plated in 10 cm dishes covered with poly-lysine coverslips at a proper concentration to let them grow at least one day. The cells are treated with 9 μM RO-3306 to block the cell cycle at the G2/M border, after 24h the cells are released to let them enter the M phase. Because not all the cells will release at the same time, starting 40 minutes from the release, 2 coverslips every 10 minutes are removed from the dish to fix the cells with 2% paraformaldehyde, this this helped accumulating a high number of mitotic cells to stain for the analysis of the mitotic spindle.

### Cell treatments

To induce replication stress and DNA damage all cell lines were treated with 2 mM Hydroxyurea for 20-24h prior to the mitotic shake-off, the four non-cancer cell lines were also treated with 20 μM RI-1, a chemical inhibitor for the DNA damage repair protein Rad51. RPE-1 cells were also treated with 1 μM and 50 μM RI-1 to test the response to different doses. To induce the depletion of CENP-A, the modified hTERT cells were treated with 0.5 μM Auxin, to activate the expression of the RNaseH1 we used 0.5 μg/ml of Doxycycline.

### Immunofluorescence

Cells grown on poly-lysine coverslips are washed three times in 1X PBS and permeabilized with 0.2% Triton diluted in 1X PBS for 5 minutes at room temperature (RT). Cells are then washed again before being blocked with PBS-0.1% Tween 20 with 5% BSA for 10 minutes. Cells are then incubated with primary antibodies for 45 minutes and washed three times with 1X PBS and 0.1% Tween 20 before being incubated with the secondary antibodies. After 30 minutes, cells are washed twice with 1X PBS and 0.1% Tween 20 and counterstained with DAPI diluted in 1X PBS. After 5 minutes, cells are washed one last time with 1X PBS and mounted using ProLong Glass Antifade mounting.

### Image acquisition

Immunofluorescence images were acquired using a Thunder Leica widefield fluorescent microscope at a 100X magnification and with a 0.2 mm z-stack.

### G1 cells Non-Random Segregation and G2 cells DNA damage analysis

After the creation of the maximum projections, the G1 couples were identified in the brightfield channels showing the cytoplasm shape around the DAPI staining of the nuclei. Only the cytoplasmic shapes surrounding two nuclei were considered for the analysis of the segregation of γH2AX. Among all the treatment conditions, all the G1 couples were divided in three categories: 1) Both cells without γH2AX: in both cells the protein is not shown; 2) Both cells with γH2AX: both cells have a visible staining of the protein; 3) NRS of γH2AX: only one of the cells shows the protein. For each condition and replicate, 50 to 70 G1 couples were scored. For the analysis of the G2 cells, the nuclei were identified thanks to the DAPI staining. Among all the treatments conditions, all the G2 cells were divided in two categories: 1) Non damaged cell: Nuclei without the presence of γH2AX; 2) Damaged cell: Nuclei with the presence of γH2AX. For each condition and replicate, 100 to 120 G2 cells were scored.

### Mitotic spindle length analysis

All the images were observed in maximum projection mode using the LAS X Office software. Each metaphase was analyzed observing 4 channels simultaneously: Blue: the DNA was stained with the DAPI to identify the correct cell cycle stage; Green: the Alexa-488 dye was used to label the tubulin; Red: the Cy3 dye was used to mark the γH2AX; Far Red: the Alexa-647 was used to stain the centromeric protein CENP-C. The metaphases were also distinguished in two categories: Non damaged metaphases: metaphases without a γH2AX signal; Damaged metaphases: metaphases with a γH2AX signal. The Draw scalebar tool was used to draw a line starting from the tip of the mitotic spindle (where all the microtubules meet) to the mid portion of the metaphase plate. After measuring the length of the mitotic spindle on both sides of the metaphases to see the overall distribution, we calculated the absolute value of the difference between the two sides to investigate how the replication stress affected the stability of chromosome segregation.

## ACKNOWLEDGMENTS

hTERT Human Fibroblasts were a kind gift of Dr. Cinzia Rinaldo (CNR Rome) (Rinaldo et al., 2012). We thank all members of the Giunta Laboratory for feedback and discussion related to this manuscript. Funding: All work in the Giunta Laboratory has been made possible thanks to support from Italian Foundation for Cancer Research - Fondazione AIRC per la Ricerca sul Cancro (AIRC Start-Up Grant 2020, Grant ID 2518 to S.G.); the Italian Ministry of Research - Ministero dell’Università e della Ricerca (MUR) through the Fondo Italiano per la Scienza (FIS2, Grant No. 2023-02742 to S.G.); Sapienza University of Rome (Ricerca Sapienza Progetti di Eccellenza, Grant No. B83C24001270005 to S.G.); and the European Research Council (ERC) under the European Union’s Horizon Europe research and innovation programme (CENTROFUN Starting Grant, Grant Agreement No. 101078838 to S.G.).

## REFERENCES

Akera, T., L. Chmátal, E. Trimm, K. Yang, C. Aonbangkhen, D. M. Chenoweth, C. Janke, R. M. Schultz, and M. A. Lampson, 2017, Spindle asymmetry drives non-Mendelian chromosome segregation: Science (New York, N.Y.), v. 358, no. 6363, p. 668–672, doi:10.1126/science.aan0092.

Altemose, N. et al., 2022, Complete genomic and epigenetic maps of human centromeres: Science, v. 376, no. 6588, p. eabl4178, doi:10.1126/science.abl4178.

Armakolas, A., and A. J. S. Klar, 2007, Left-Right Dynein Motor Implicated in Selective Chromatid Segregation in Mouse Cells: Science, v. 315, no. 5808, p. 100–101, doi:10.1126/science.1129429.

Arnaudeau, C., C. Lundin, and T. Helleday, 2001, DNA double-strand breaks associated with replication forks are predominantly repaired by homologous recombination involving an exchange mechanism in mammalian cells: Journal of Molecular Biology, v. 307, no. 5, p. 1235– 1245, doi:10.1006/jmbi.2001.4564.

Balzano, E., F. Pelliccia, and S. Giunta, 2021, Genome (in)stability at tandem repeats: Seminars in Cell C Developmental Biology, v. 113, p. 97–112, doi:10.1016/j.semcdb.2020.10.003.

Barra, V., and D. Fachinetti, 2018, The dark side of centromeres: types, causes and consequences of structural abnormalities implicating centromeric DNA, 1: Nature Communications, v. 9, no. 1, p. 4340, doi:10.1038/s41467-018-06545-y.

Bhat, K. P., and D. Cortez, 2018, RPA and RAD51: fork reversal, fork protection, and genome stability: Nature Structural C Molecular Biology, v. 25, no. 6, p. 446–453, doi:10.1038/s41594-018-0075-z.

Bhowmick, R., M. Lerdrup, S. A. Gadi, G. G. Rossetti, M. I. Singh, Y. Liu, T. D. Halazonetis, and I. D. Hickson, 2022, RAD51 protects human cells from transcription-replication conflicts: Molecular Cell, v. 82, no. 18, p. 3366–3381.e9, doi:10.1016/j.molcel.2022.07.010.

Birchler, J. A., Z. Gao, and F. Han, 2009, A tale of two centromeres—diversity of structure but conservation of function in plants and animals: Functional & Integrative Genomics, v. 9, no. 1, p. 7–13, doi:10.1007/s10142-008-0104-9.

Black, E. M., and S. Giunta, 2018, Repetitive Fragile Sites: Centromere Satellite DNA as a Source of Genome Instability in Human Diseases, 12: Genes, v. 9, no. 12, p. 615, doi:10.3390/genes9120615.

Blower, M. D., B. A. Sullivan, and G. H. Karpen, 2002, Conserved Organization of Centromeric Chromatin in Flies and Humans: Developmental Cell, v. 2, no. 3, p. 319–330, doi:10.1016/S1534-5807(02)00135-1.

Bosco, N. et al., 2023, KaryoCreate: A CRISPR-based technology to study chromosome-specific aneuploidy by targeting human centromeres: Cell, p. S0092–8674(23)00326–4, doi:10.1016/j.cell.2023.03.029.

Bosco, N., F. Pelliccia, and A. Rocchi, 2010, Characterization of FRA7B, a human common fragile site mapped at the 7p chromosome terminal region: Cancer Genetics and Cytogenetics, v. 202, no. 1, p. 47–52, doi:10.1016/j.cancergencyto.2010.06.008.

Boteva, L., R.-S. Nozawa, C. Naughton, K. Samejima, W. C. Earnshaw, and N. Gilbert, 2020, Common Fragile Sites Are Characterized by Faulty Condensin Loading after Replication Stress: Cell Reports, v. 32, no. 12, p. 108177, doi:10.1016/j.celrep.2020.108177.

Budke, B., H. L. Logan, J. H. Kalin, A. S. Zelivianskaia, W. Cameron McGuire, L. L. Miller, J. M. Stark, A. P. Kozikowski, D. K. Bishop, and P. P. Connell, 2012, RI-1: a chemical inhibitor of RAD51 that disrupts homologous recombination in human cells: Nucleic Acids Research, v. 40, no. 15, p. 7347–7357, doi:10.1093/nar/gks353.

Cairns, J., 1975, Mutation selection and the natural history of cancer: Nature, v. 255, no. 5505, p. 197– 200, doi:10.1038/255197a0.

Celeste, A. et al., 2002, Genomic instability in mice lacking histone H2AX: Science, v. 296, no. 5569, p. 922–927, doi:10.1126/science.1069398.

Celeste, A., O. Fernandez-Capetillo, M. J. Kruhlak, D. R. Pilch, D. W. Staudt, A. Lee, R. F. Bonner, W. M. Bonner, and A. Nussenzweig, 2003, Histone H2AX phosphorylation is dispensable for the initial recognition of DNA breaks: Nature Cell Biology, v. 5, no. 7, p. 675–679, doi:10.1038/ncb1004.

Chan, F. L., O. J. Marshall, R. Saffery, B. W. Kim, E. Earle, K. H. A. Choo, and L. H. Wong, 2012, Active transcription and essential role of RNA polymerase II at the centromere during mitosis: Proceedings of the National Academy of Sciences of the United States of America, v. 109, no. 6, p. 1979–1984, doi:10.1073/pnas.1108705109.

Chan, F. L., and L. H. Wong, 2012, Transcription in the maintenance of centromere chromatin identity: Nucleic Acids Research, v. 40, no. 22, p. 11178–11188, doi:10.1093/nar/gks921.

Charville, G. W., and T. A. Rando, 2013, The Mortal Strand Hypothesis: Non-random chromosome inheritance and the biased segregation of damaged DNA: Seminars in cell C developmental biology, v. 24, no. 0, p. 10.1016/j.semcdb.2013.05.006, doi:10.1016/j.semcdb.2013.05.006.

Chittoor, S. S., and S. Giunta, 2024, Comparative analysis of predicted DNA secondary structures infers complex human centromere topology: American Journal of Human Genetics, v. 111, no. 12, p. 2707–2719, doi:10.1016/j.ajhg.2024.10.016.

Conboy, M. J., A. O. Karasov, and T. A. Rando, 2007, High Incidence of Non-Random Template Strand Segregation and Asymmetric Fate Determination In Dividing Stem Cells and their Progeny: PLOS Biology, v. 5, no. 5, p. e102, doi:10.1371/journal.pbio.0050102.

Dalal, Y., T. Furuyama, D. Vermaak, and S. Henikoff, 2007, Structure, dynamics, and evolution of centromeric nucleosomes: Proceedings of the National Academy of Sciences, v. 104, no. 41, p. 15974–15981, doi:10.1073/pnas.0707648104.

Debatisse, M., B. Le Tallec, A. Letessier, B. Dutrillaux, and O. Brison, 2012, Common fragile sites: mechanisms of instability revisited: Trends in Genetics, v. 28, no. 1, p. 22–32, doi:10.1016/j.tig.2011.10.003.

Di Tommaso, E., and S. Giunta, 2023, Dynamic interplay between human alpha-satellite DNA structure and centromere functions: Seminars in Cell C Developmental Biology, doi:10.1016/j.semcdb.2023.10.002.

Di Tommaso, E., V. de Turris, P. Choppakatla, H. Funabiki, and S. Giunta, 2023, Visualization of the three-dimensional structure of the human centromere in mitotic chromosomes by superresolution microscopy: Molecular Biology of the Cell, v. 34, no. 6, p. ar61, doi:10.1091/mbc.E22-08-0332.

Dumont, M. et al., 2020, Human chromosome-specific aneuploidy is influenced by DNA-dependent centromeric features: The EMBO journal, v. 39, no. 2, p. e102924, doi:10.15252/embj.2019102924.

El Achkar, E., M. Gerbault-Seureau, M. Muleris, B. Dutrillaux, and M. Debatisse, 2005, Premature condensation induces breaks at the interface of early and late replicating chromosome bands bearing common fragile sites: Proceedings of the National Academy of Sciences of the United States of America, v. 102, no. 50, p. 18069–18074, doi:10.1073/pnas.0506497102.

Esashi, F., V. E. Galkin, X. Yu, E. H. Egelman, and S. C. West, 2007, Stabilization of RAD51 nucleoprotein filaments by the C-terminal region of BRCA2, 6: Nature Structural C Molecular Biology, v. 14, no. 6, p. 468–474, doi:10.1038/nsmb1245.

Ferrand, J., J. Dabin, O. Chevallier, M. Kane-Charvin, A. Kupai, J. Hrit, S. B. Rothbart, and S. E. Polo, 2025, Mitotic chromatin marking governs the segregation of DNA damage: Nature Communications, v. 16, no. 1, p. 746, doi:10.1038/s41467-025-56090-8.

Ginda, K., I. Santi, D. Bousbaine, J. Zakrzewska-Czerwińska, D. Jakimowicz, and J. McKinney, 2017, The studies of ParA and ParB dynamics reveal asymmetry of chromosome segregation in mycobacteria: Molecular Microbiology, v. 105, no. 3, p. 453–468, doi:10.1111/mmi.13712.

Giunta, S. et al., 2021, CENP-A chromatin prevents replication stress at centromeres to avoid structural aneuploidy: Proceedings of the National Academy of Sciences, v. 118, no. 10, p. e2015634118, doi:10.1073/pnas.2015634118.

Giunta, S., and H. Funabiki, 2017, Integrity of the human centromere DNA repeats is protected by CENP-A, CENP-C, and CENP-T: Proceedings of the National Academy of Sciences, v. 114, no. 8, p. 1928–1933, doi:10.1073/pnas.1615133114.

Glover, T. W., C. Berger, J. Coyle, and B. Echo, 1984, DNA polymerase α inhibition by aphidicolin induces gaps and breaks at common fragile sites in human chromosomes: Human Genetics, v. 67, no. 2, p. 136–142, doi:10.1007/BF00272988.

Gordon, D. J., B. Resio, and D. Pellman, 2012, Causes and consequences of aneuploidy in cancer: Nature Reviews Genetics, v. 13, no. 3, p. 189–203, doi:10.1038/nrg3123.

Hayden, K. E., 2012, Human centromere genomics: now it’s personal: Chromosome Research, v. 20, no. 5, p. 621–633, doi:10.1007/s10577-012-9295-y.

Heller, R. C., and K. J. Marians, 2006, Replisome assembly and the direct restart of stalled replication forks: Nature Reviews. Molecular Cell Biology, v. 7, no. 12, p. 932–943, doi:10.1038/nrm2058.

Helmrich, A., M. Ballarino, and L. Tora, 2011, Collisions between replication and transcription complexes cause common fragile site instability at the longest human genes: Molecular Cell, v. 44, no. 6, p. 966–977, doi:10.1016/j.molcel.2011.10.013.

Kabeche, L., H. D. Nguyen, R. Buisson, and L. Zou, 2018, A mitosis-specific and R loop-driven ATR pathway promotes faithful chromosome segregation: Science (New York, N.Y.), v. 359, no. 6371, p. 108–114, doi:10.1126/science.aan6490.

Karpowicz, P., C. Morshead, A. Kam, E. Jervis, J. Ramunas, V. Cheng, and D. van der Kooy, 2005, Support for the immortal strand hypothesis: neural stem cells partition DNA asymmetrically in vitro: Journal of Cell Biology, v. 170, no. 5, p. 721–732, doi:10.1083/jcb.200502073.

Kasinathan, S., and S. Henikoff, 2018, Non-B-Form DNA Is Enriched at Centromeres: Molecular Biology and Evolution, v. 35, no. 4, p. 949–962, doi:10.1093/molbev/msy010.

Le Beau, M. M., F. V. Rassool, M. E. Neilly, R. Espinosa III, T. W. Glover, D. I. Smith, and T. W. McKeithan, 1998, Replication of a Common Fragile Site, FRA3B, Occurs Late in S Phase and is Delayed Further Upon Induction: Implications for the Mechanism of Fragile Site Induction: Human Molecular Genetics, v. 7, no. 4, p. 755–761, doi:10.1093/hmg/7.4.755.

Li, X., and W.-D. Heyer, 2008, Homologous recombination in DNA repair and DNA damage tolerance: Cell Research, v. 18, no. 1, p. 99–113, doi:10.1038/cr.2008.1.

Liu, D., and M. A. Lampson, 2009, Regulation of kinetochore–microtubule attachments by Aurora B kinase: Biochemical Society Transactions, v. 37, no. 5, p. 976–980, doi:10.1042/BST0370976.

Lundin, C., K. Erixon, C. Arnaudeau, N. Schultz, D. Jenssen, M. Meuth, and T. Helleday, 2002, Different roles for nonhomologous end joining and homologous recombination following replication arrest in mammalian cells: Molecular and Cellular Biology, v. 22, no. 16, p. 5869–5878, doi:10.1128/MCB.22.16.5869-5878.2002.

Maccaroni, K., E. Balzano, F. Mirimao, S. Giunta, and F. Pelliccia, 2020, Impaired Replication Timing Promotes Tissue-Specific Expression of Common Fragile Sites: Genes, v. 11, no. 3, p. 326, doi:10.3390/genes11030326.

Mailand, N., I. Gibbs-Seymour, and S. Bekker-Jensen, 2013, Regulation of PCNA-protein interactions for genome stability: Nature Reviews. Molecular Cell Biology, v. 14, no. 5, p. 269–282, doi:10.1038/nrm3562.

Marnef, A., and G. Legube, 2021, R-loops as Janus-faced modulators of DNA repair: Nature Cell Biology, v. 23, no. 4, p. 305–313, doi:10.1038/s41556-021-00663-4.

Massey, D. J., D. Kim, K. E. Brooks, M. B. Smolka, and A. Koren, 2019, Next-Generation Sequencing Enables Spatiotemporal Resolution of Human Centromere Replication Timing: Genes, v. 10, no. 4, p. 269, doi:10.3390/genes10040269.

Mays, J. C., S. Mei, N. Bosco, X. Zhao, J. J. Bianchi, G. R. Kidiyoor, L. J. Holt, and T. Davoli, 2023, KaryoTap Enables Aneuploidy Detection in Thousands of Single Human Cells: bioRxiv, p. 2023.09.08.555746, doi:10.1101/2023.09.08.555746.

McKinley, K. L., and I. M. Cheeseman, 2016, The molecular basis for centromere identity and function, 1: Nature Reviews Molecular Cell Biology, v. 17, no. 1, p. 16–29, doi:10.1038/nrm.2015.5.

McNulty, S. M., L. L. Sullivan, and B. A. Sullivan, 2017, Human Centromeres Produce Chromosome-Specific and Array-Specific Alpha Satellite Transcripts that Are Complexed with CENP-A and CENP-C: Developmental Cell, v. 42, no. 3, p. 226–240.e6, doi:10.1016/j.devcel.2017.07.001.

Musacchio, A., 2015, The Molecular Biology of Spindle Assembly Checkpoint Signaling Dynamics: Current Biology, v. 25, no. 20, p. R1002–R1018, doi:10.1016/j.cub.2015.08.051.

Ortega, P., J. A. Mérida-Cerro, A. G. Rondón, B. Gómez-González, and A. Aguilera, 2021, DNA-RNA hybrids at DSBs interfere with repair by homologous recombination: eLife, v. 10, p. e69881, doi:10.7554/eLife.69881.

Papini, D., M. D. Levasseur, and J. M. G. Higgins, 2021, The Aurora B gradient sustains kinetochore stability in anaphase: Cell Reports, v. 37, no. 6, doi:10.1016/j.celrep.2021.109818.

Petermann, E., M. L. Orta, N. Issaeva, N. Schultz, and T. Helleday, 2010, Hydroxyurea-stalled replication forks become progressively inactivated and require two different RAD51-mediated pathways for restart and repair: Molecular Cell, v. 37, no. 4, p. 492–502, doi:10.1016/j.molcel.2010.01.021.

Petryk, N., M. Dalby, A. Wenger, C. B. Stromme, A. Strandsby, R. Andersson, and A. Groth, 2018, MCM2 promotes symmetric inheritance of modified histones during DNA replication: Science, v. 361, no. 6409, p. 1389–1392, doi:10.1126/science.aau0294.

Potten, C. S., G. Owen, and D. Booth, 2002, Intestinal stem cells protect their genome by selective segregation of template DNA strands: Journal of Cell Science, v. 115, no. 11, p. 2381–2388, doi:10.1242/jcs.115.11.2381.

Quinet, A., D. Lemaçon, and A. Vindigni, 2017, Replication fork reversal: players and guardians: Molecular cell, v. 68, no. 5, p. 830–833, doi:10.1016/j.molcel.2017.11.022.

Ribeiro, S. A., P. Vagnarelli, Y. Dong, T. Hori, B. F. McEwen, T. Fukagawa, C. Flors, and W. C. Earnshaw, 2010, A super-resolution map of the vertebrate kinetochore: Proceedings of the National Academy of Sciences, v. 107, no. 23, p. 10484–10489, doi:10.1073/pnas.1002325107.

Rinaldo, C. et al., 2012, HIPK2 Controls Cytokinesis and Prevents Tetraploidization by Phosphorylating Histone H2B at the Midbody: Molecular Cell, v. 47, no. 1, p. 87–98, doi:10.1016/j.molcel.2012.04.029.

Saayman, X., E. Graham, W. J. Nathan, A. Nussenzweig, and F. Esashi, 2023, Centromeres as universal hotspots of DNA breakage, driving RAD51-mediated recombination during quiescence: Molecular Cell, v. 83, no. 4, p. 523–538.e7, doi:10.1016/j.molcel.2023.01.004.

Saintigny, Y., F. Delacôte, G. Varès, F. Petitot, S. Lambert, D. Averbeck, and B. S. Lopez, 2001, Characterization of homologous recombination induced by replication inhibition in mammalian cells: The EMBO journal, v. 20, no. 14, p. 3861–3870, doi:10.1093/emboj/20.14.3861.

Santaguida, S., and A. Amon, 2015, Short- and long-term effects of chromosome mis-segregation and aneuploidy: Nature Reviews Molecular Cell Biology, v. 16, no. 8, p. 473–485, doi:10.1038/nrm4025.

Santaguida, S., and A. Musacchio, 2009, The life and miracles of kinetochores: The EMBO Journal, v. 28, no. 17, p. 2511–2531, doi:10.1038/emboj.2009.173.

Scelfo, A. et al., 2024, Specialized replication mechanisms maintain genome stability at human centromeres: Molecular Cell, v. 84, no. 6, p. 1003–1020.e10, doi:10.1016/j.molcel.2024.01.018.

Schalch, T., and F. A. Steiner, 2017, Structure of centromere chromatin: from nucleosome to chromosomal architecture: Chromosoma, v. 126, no. 4, p. 443–455, doi:10.1007/s00412-016-0620-7.

Schueler, M. G., and B. A. Sullivan, 2006, Structural and Functional Dynamics of Human Centromeric Chromatin: Annual Review of Genomics and Human Genetics, v. 7, no. 1, p. 301–313, doi:10.1146/annurev.genom.7.080505.115613.

Shinin, V., B. Gayraud-Morel, D. Gomès, and S. Tajbakhsh, 2006, Asymmetric division and cosegregation of template DNA strands in adult muscle satellite cells: Nature Cell Biology, v. 8, no. 7, p. 677–682, doi:10.1038/ncb1425.

Smith, G. H., 2005, Label-retaining epithelial cells in mouse mammary gland divide asymmetrically and retain their template DNA strands: Development, v. 132, no. 4, p. 681–687, doi:10.1242/dev.01609.

Smith, D. I., Y. Zhu, S. McAvoy, and R. Kuhn, 2006, Common fragile sites, extremely large genes, neural development and cancer: Cancer Letters, v. 232, no. 1, p. 48–57, doi:10.1016/j.canlet.2005.06.049.

Stern, B. M., and A. W. Murray, 2001, Lack of tension at kinetochores activates the spindle checkpoint in budding yeast: Current Biology, v. 11, no. 18, p. 1462–1467, doi:10.1016/S0960-9822(01)00451-1.

Sullivan, B. A., and G. H. Karpen, 2004, Centromeric chromatin exhibits a histone modification pattern that is distinct from both euchromatin and heterochromatin, 11: Nature Structural C Molecular Biology, v. 11, no. 11, p. 1076–1083, doi:10.1038/nsmb845.

Ten Hagen, K. G., D. M. Gilbert, H. F. Willard, and S. N. Cohen, 1990, Replication timing of DNA sequences associated with human centromeres and telomeres: Molecular and Cellular Biology, v. 10, no. 12, p. 6348–6355, doi:10.1128/mcb.10.12.6348-6355.1990.

Valdiglesias, V., S. Giunta, M. Fenech, M. Neri, and S. Bonassi, 2013, γH2AX as a marker of DNA double strand breaks and genomic instability in human population studies: Mutation Research, v. 753, no. 1, p. 24–40, doi:10.1016/j.mrrev.2013.02.001.

Wenger, A. et al., 2023, Symmetric inheritance of parental histones governs epigenome maintenance and embryonic stem cell identity: Nature Genetics, v. 55, no. 9, p. 1567–1578, doi:10.1038/s41588-023-01476-x.

Willard, H. F., 1985, Chromosome-specific organization of human alpha satellite DNA: American Journal of Human Genetics, v. 37, no. 3, p. 524–532.

Willard, H. F., and J. S. Waye, 1987, Hierarchical order in chromosome-specific human alpha satellite DNA: Trends in Genetics, v. 3, p. 192–198, doi:10.1016/0168-9525(87)90232-0.

Xing, M. et al., 2020, Replication Stress Induces ATR/CHK1-Dependent Nonrandom Segregation of Damaged Chromosomes: Molecular Cell, v. 78, no. 4, p. 714–724.e5, doi:10.1016/j.molcel.2020.04.005.

Yadlapalli, S., and Y. M. Yamashita, 2013, Chromosome-specific nonrandom sister chromatid segregation during stem-cell division: Nature, v. 498, no. 7453, p. 251–254, doi:10.1038/nature12106.

Yeh, E., J. Haase, L. V. Paliulis, A. Joglekar, L. Bond, D. Bouck, E. D. Salmon, and K. S. Bloom, 2008, Pericentric Chromatin Is Organized into an Intramolecular Loop in Mitosis: Current Biology, v. 18, no. 2, p. 81–90, doi:10.1016/j.cub.2007.12.019.

Yilmaz, D., A. Furst, K. Meaburn, A. Lezaja, Y. Wen, M. Altmeyer, B. Reina-San-Martin, and E. Soutoglou, 2021, Activation of homologous recombination in G1 preserves centromeric integrity, 7890: Nature, v. 600, no. 7890, p. 748–753, doi:10.1038/s41586-021-04200-z.

Zeman, M. K., and K. A. Cimprich, 2014, Causes and consequences of replication stress: Nature Cell Biology, v. 16, no. 1, p. 2–9, doi:10.1038/ncb2897.

Zinkowski, R. P., J. Meyne, and B. R. Brinkley, 1991, The centromere-kinetochore complex: a repeat subunit model.: Journal of Cell Biology, v. 113, no. 5, p. 1091–1110, doi:10.1083/jcb.113.5.1091.

