## Supplementary Figures for "Replication stress at centromeres biases the segregation of DNA damage"

### Replication stress at centromeres contributes to Non-Random Segregation of DNA damage

Supplementary Figures and Figure Legends

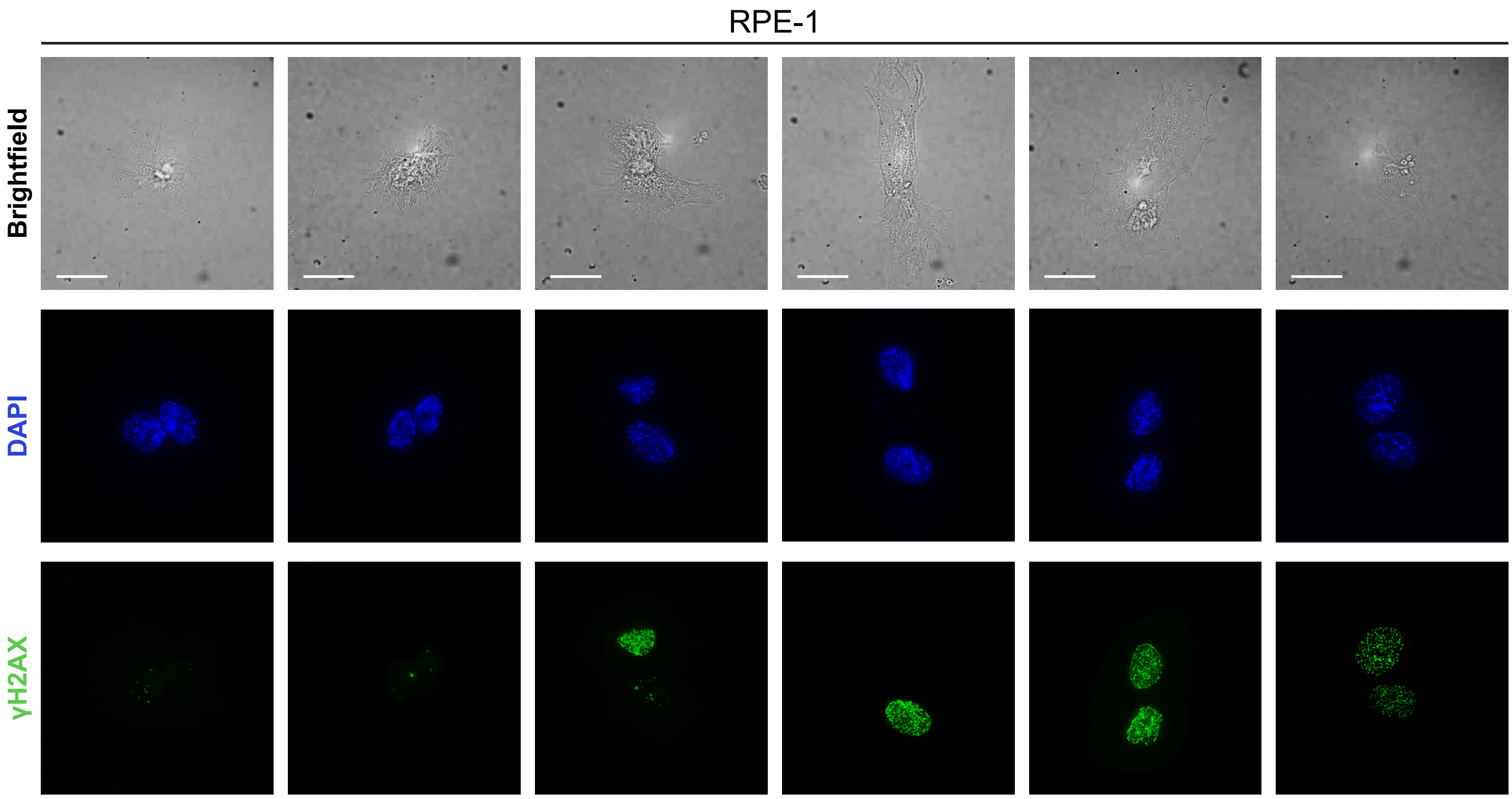
RPE-1
Brightfield
DAPI
γH2AX

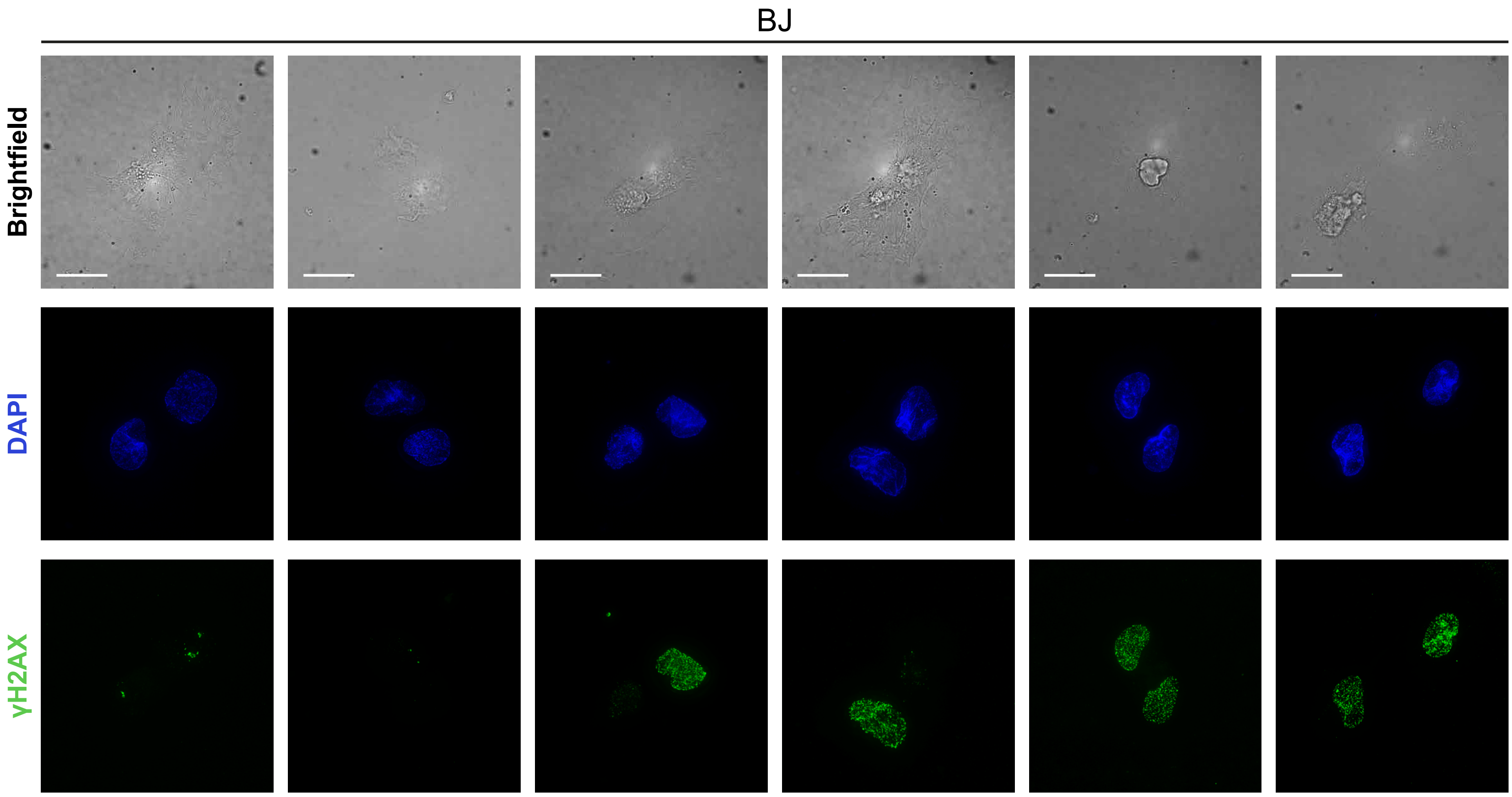
BJ
Brightfield
DAPI
γH2AX

**Supplementary Figure 1: Segregation patterns of γH2AX in RPE-1 and BJ cells.** Examples of RPE-1 and BJ cells showing: both G1 without DNA damage, NRS of DNA damage, both G1 with DNA damage, scalebar is 20 μm.

hCEC D29

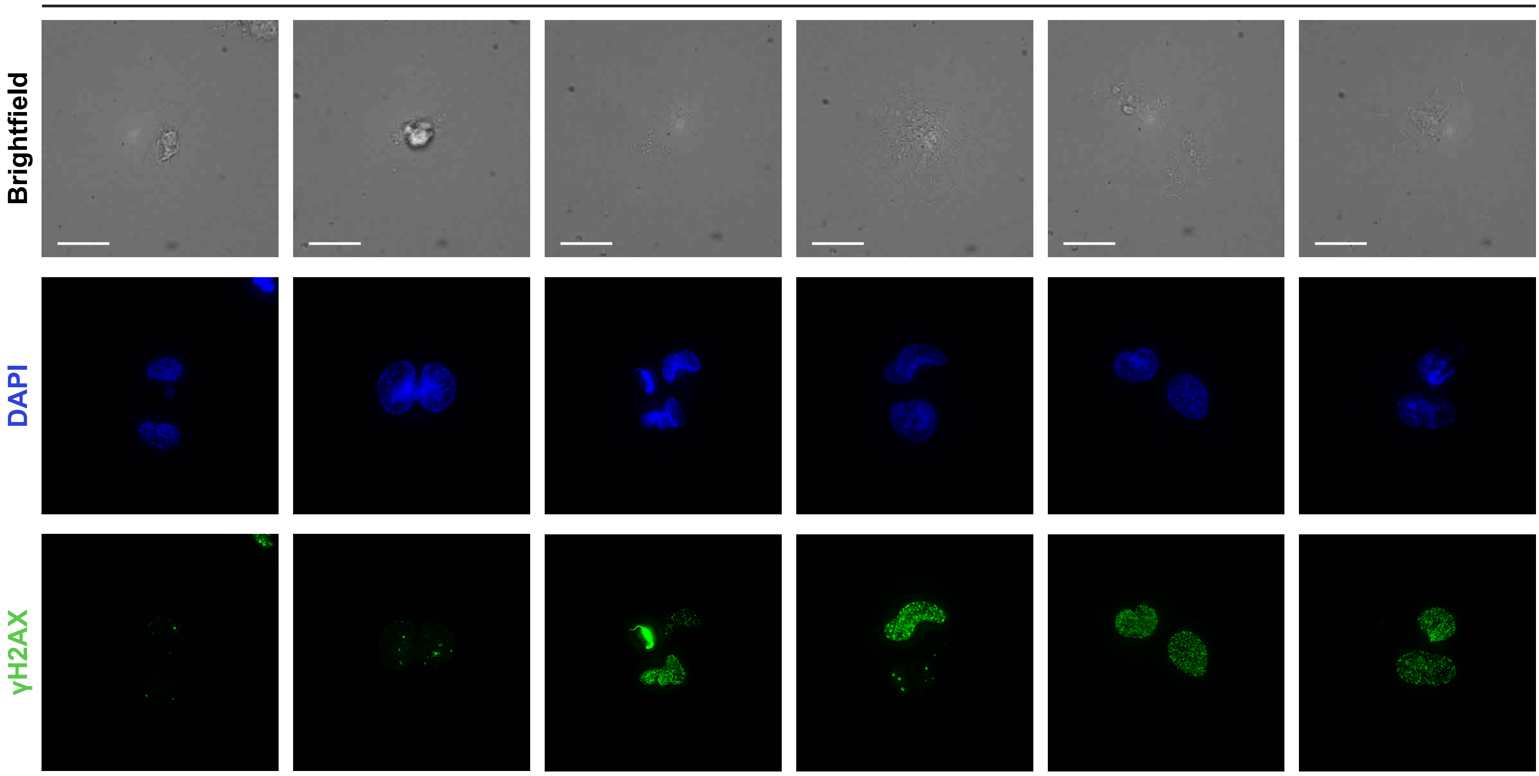


Human Fibroblasts

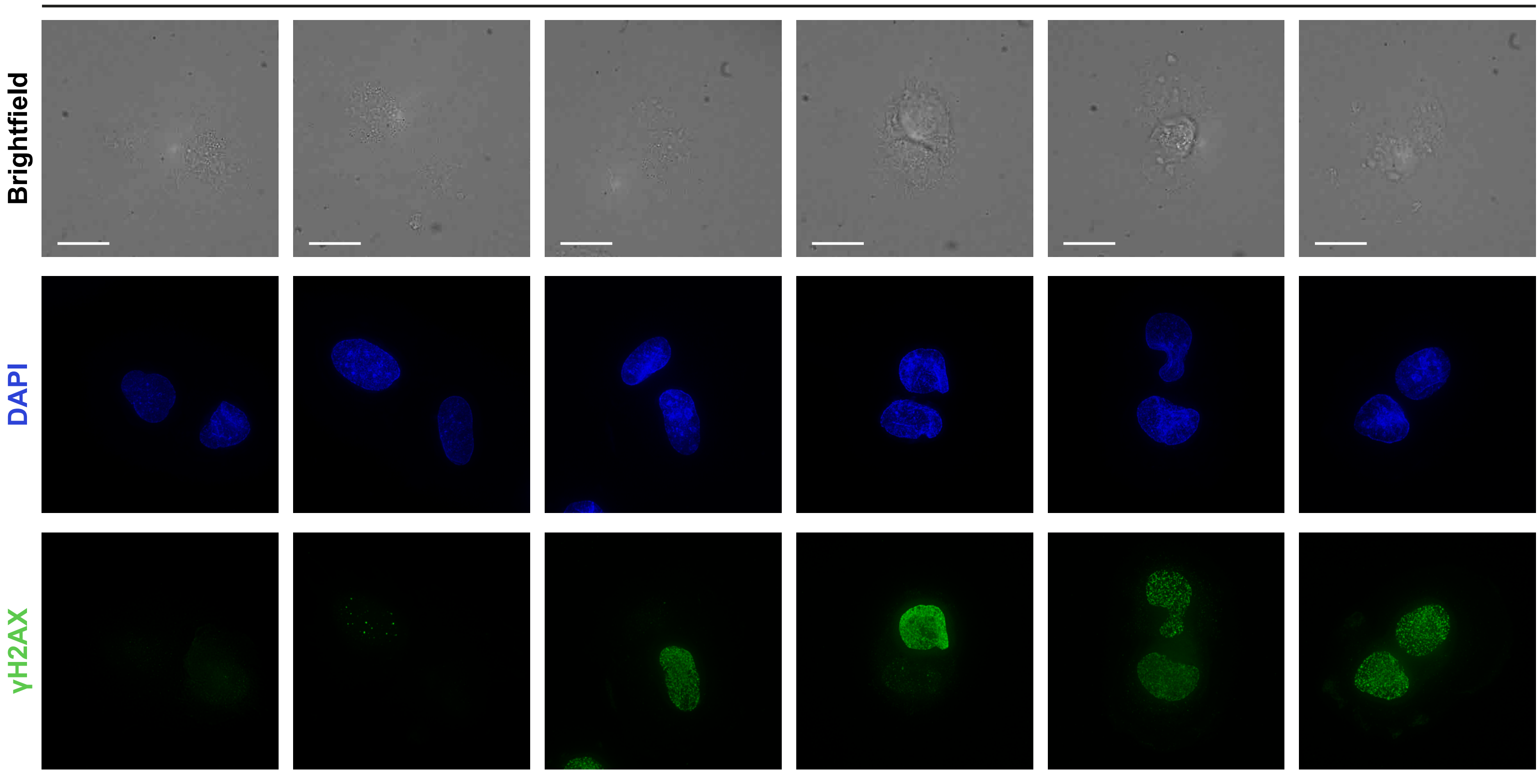

**Supplementary Figure 2: Segregation patterns of γH2AX in hCEC D29, and Human Fibroblasts cells.** Examples of hCEC D29 and Human Fibroblasts cells showing: both G1 without DNA damage, NRS of DNA damage, both G1 with DNA damage, scalebar is 20 μm.

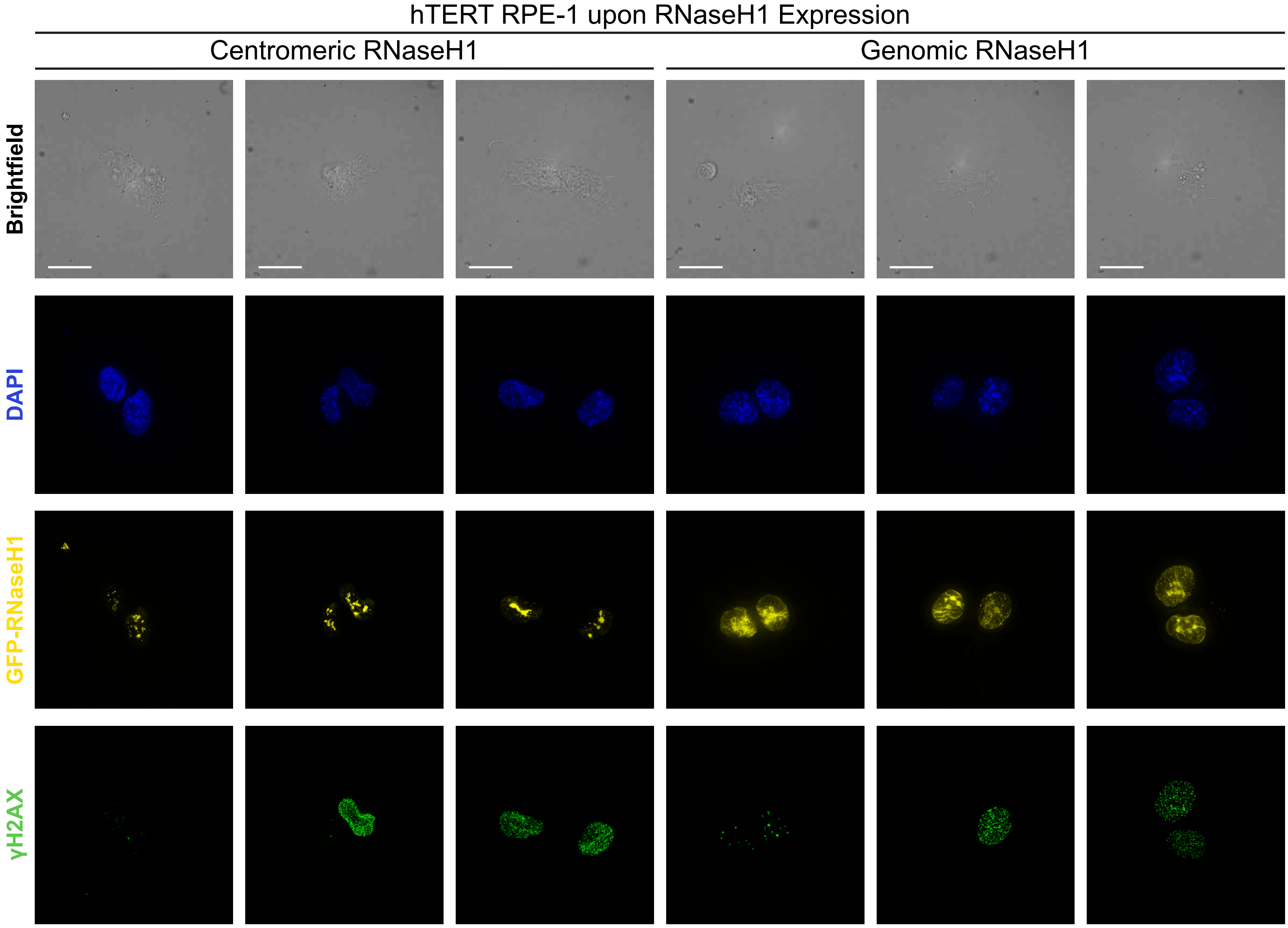
hTERT RPE-1 upon RNaseH1 Expression
Centromeric RNaseH1
Genomic RNaseH1
Brightfield
DAPI
GFP-RNaseH1
γH2AX

**Supplementary Figure 3: Visualization of RNaseH1 expression in hTERT RPE-1 cells.** Examples of hTERT RPE-1 cells showing the Doxycycline inducible expression of RNaseH1 fused with GFP, highlighting the different localization between the Centromeric and Genomic one.

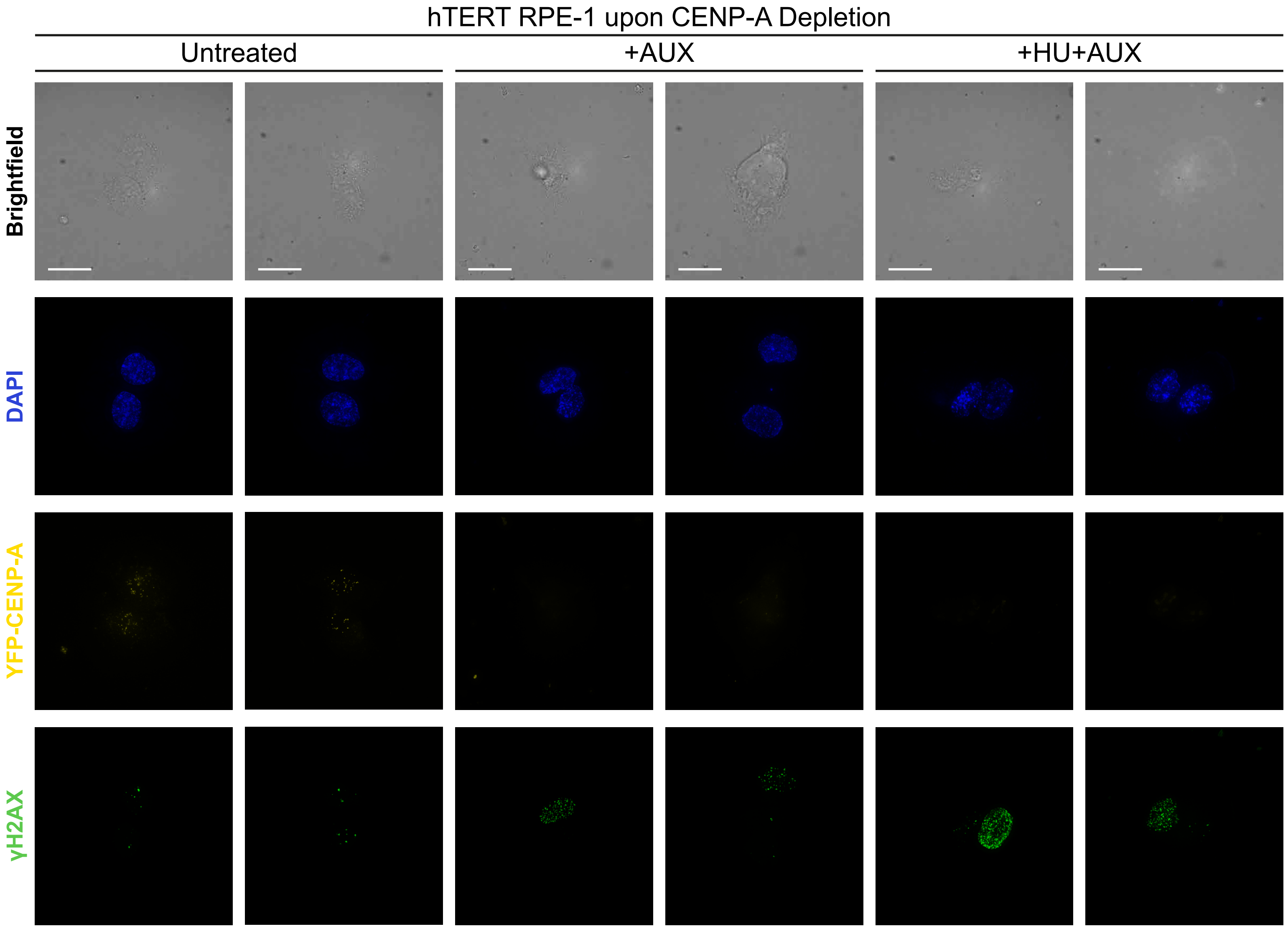
hTERT RPE-1 upon CENP-A Depletion
Untreated
+AUX
+HU+AUX
Brightfield
DAPI
YFP-CENP-A
γH2AX

**Supplementary Figure 4: Visualization of CENP-A depletion in hTERT RPE-1 cells.** Examples of hTERT RPE-1 cells showing CENP-A fused to YFP being depleted after Auxin treatment.

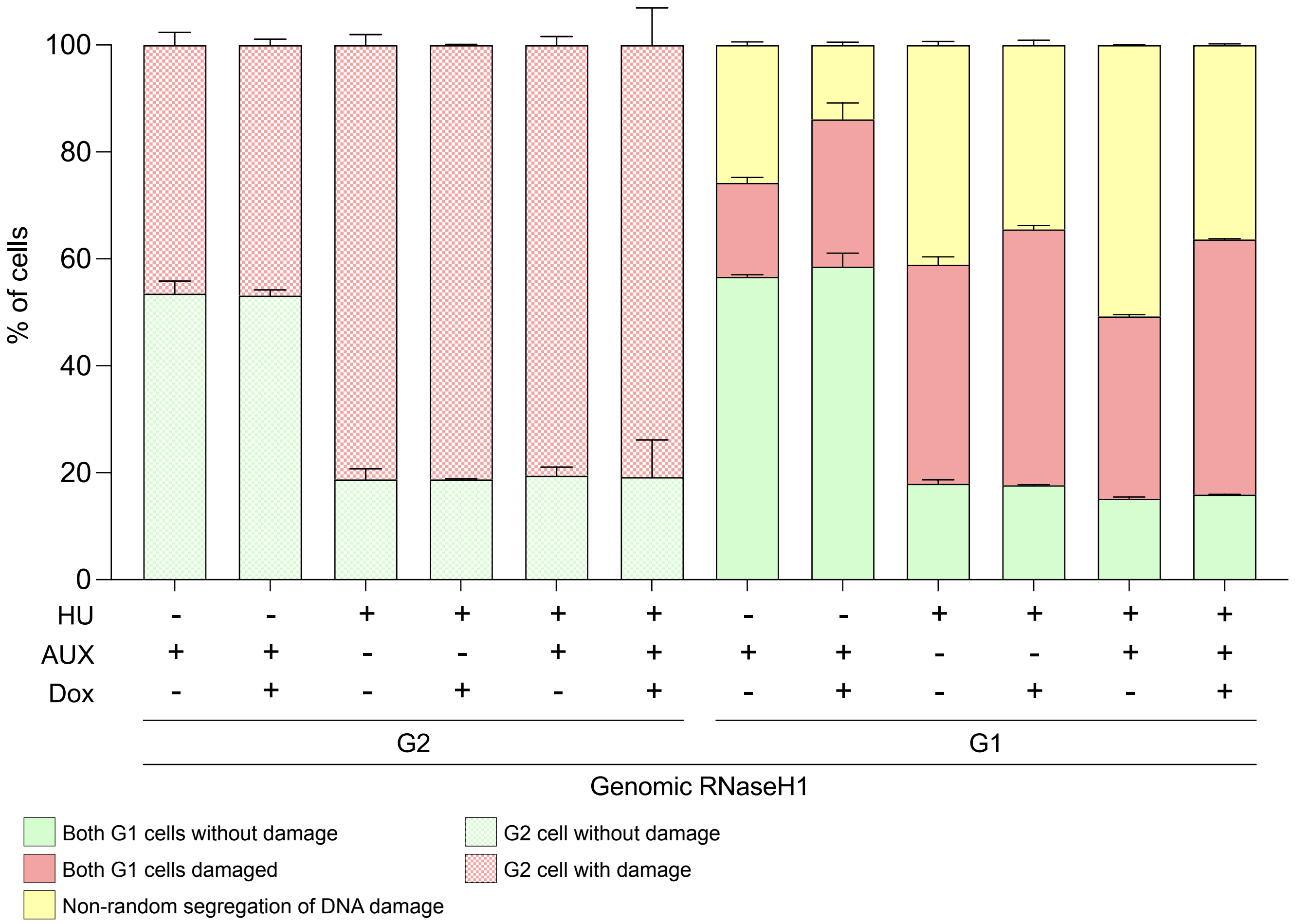

% of cells
100
80
60
40
20
0
HU
AUX
Dox
G2
G1
Genomic RNaseH1
Both G1 cells without damage
Both G1 cells damaged
Non-random segregation of DNA damage
G2 cell without damage
G2 cell with damage

**Supplementary Figure 5: Combinatorial effects of CENP-A depletion and R-loops removal on replication stress and NRS.** On the left side of the graph, quantification of the genomic RNaseH1 RPE-1 G2 cells as the examples in Fig. 3C, with and without HU, AUX and Dox treatments. For each individual experiment, 100-120 G2s were counted. On the right side of the graph, quantification of the genomic RNaseH1 RPE-1 G1 cells as the examples in Fig. 3C, with and without HU, AUX and Dox treatments. For each individual experiment, 55-80 G1 couples were counted. The error bars show the SD of the mean across three biological replicates.

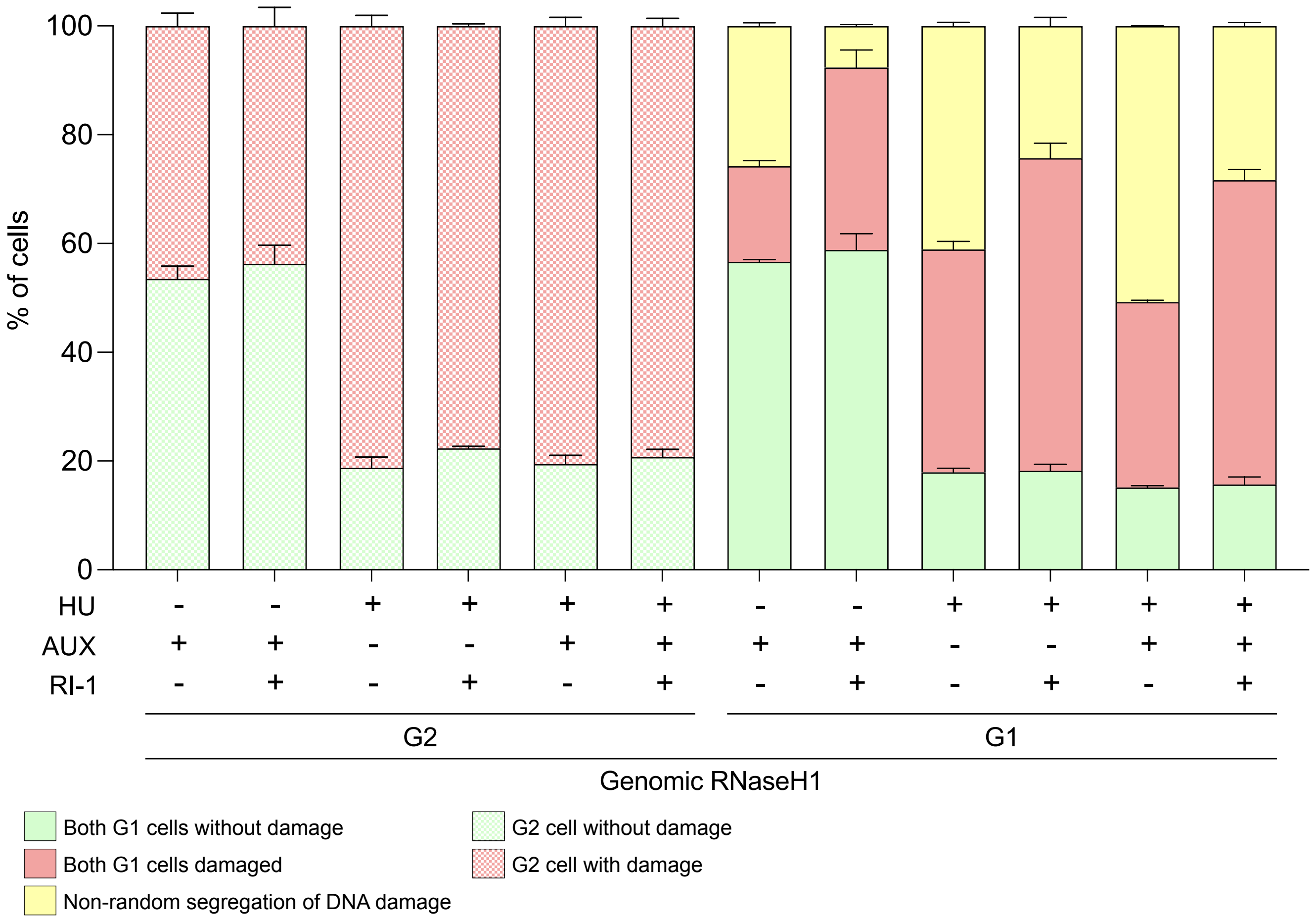

100
80
60
40
20
0
% of cells
HU
AUX
RI-1
G2
G1
Genomic RNaseH1
Both G1 cells without damage
Both G1 cells damaged
Non-random segregation of DNA damage
G2 cell without damage
G2 cell with damage

**Supplementary Figure 6: Combinatorial effects of CENP-A depletion and Rad51 inhibition on replication stress and NRS.** On the left side of the graph, quantification of the genomic RNaseH1 RPE-1 G2 cells as the examples in Fig. 3C, with and without HU, AUX and RI-1 treatments. For each individual experiment, 100-120 G2s were counted. On the right side of the graph, quantification of the genomic RNaseH1 RPE-1 G1 cells as the examples in Fig. 3C, with and without HU, AUX and RI-1 treatments. For each individual experiment, 55-80 G1 couples were counted. The error bars show the SD of the mean across three biological replicates

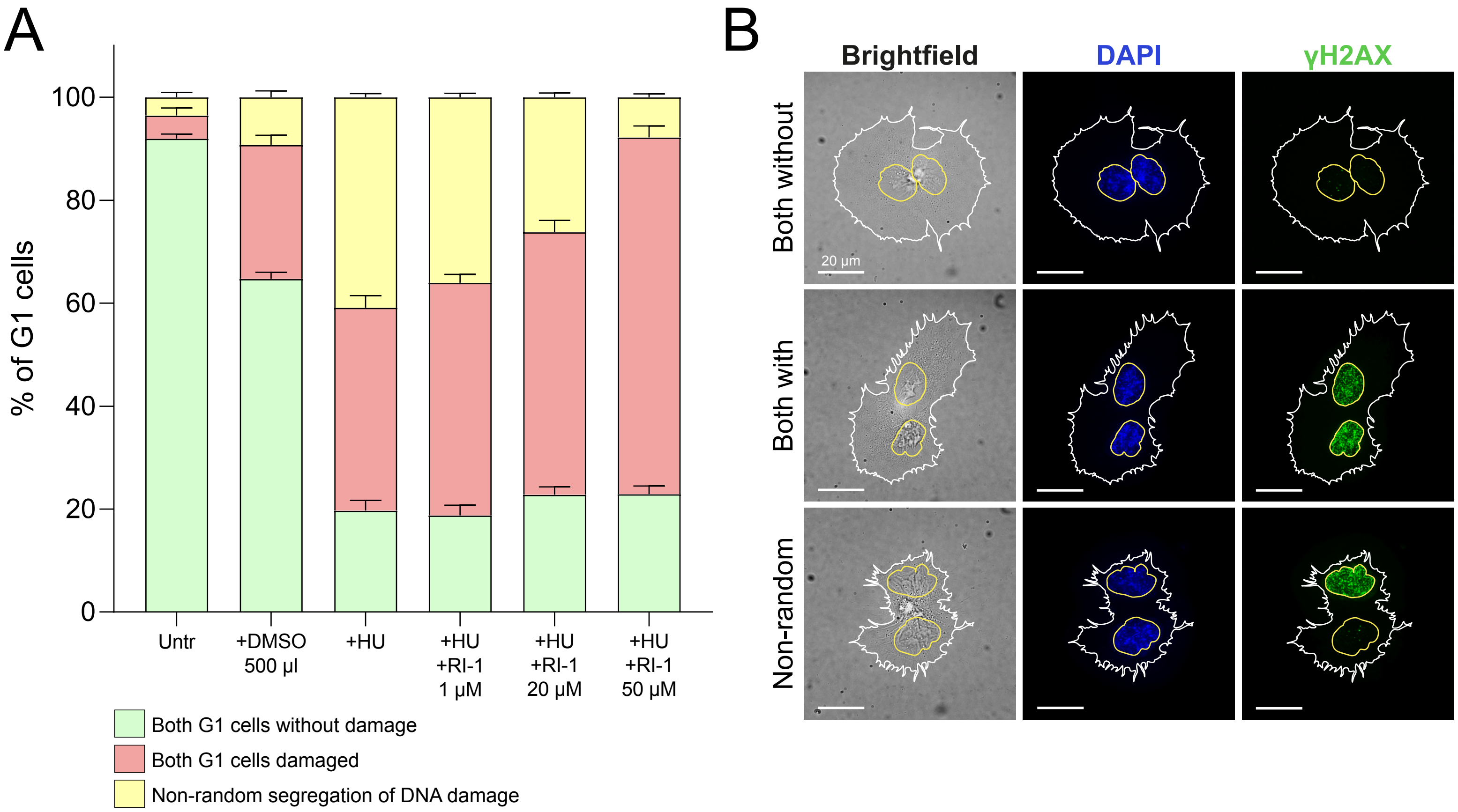
A
% of G1 cells
0
20
40
60
80
100
Untr
+DMSO 500 µl
+HU
+HU +RI-1 1 µM
+HU +RI-1 20 µM
+HU +RI-1 50 µM
Both G1 cells without damage
Both G1 cells damaged
Non-random segregation of DNA damage
B
Brightfield
DAPI
γH2AX
Both without
Both with
Non-random
20 µm

**Supplementary Figure 7: Rad51 inhibition is dependent on the doses used.** (A) Quantification of the RPE-1 G1 cells as the examples in B, with and without HU treatment, and with different doses of RI-1. For each individual experiment, 55-80 G1 couples were counted. The error bars show the SD of the mean across three biological replicates. (B) Examples of RPE-1 and BJ cells showing: both G1 without DNA damage, both G1 with DNA damage, NRS of DNA damage. The white outline highlights the cytoplasm shape, while the yellow one highlights the nucleus shape, scalebar is 20 μm.

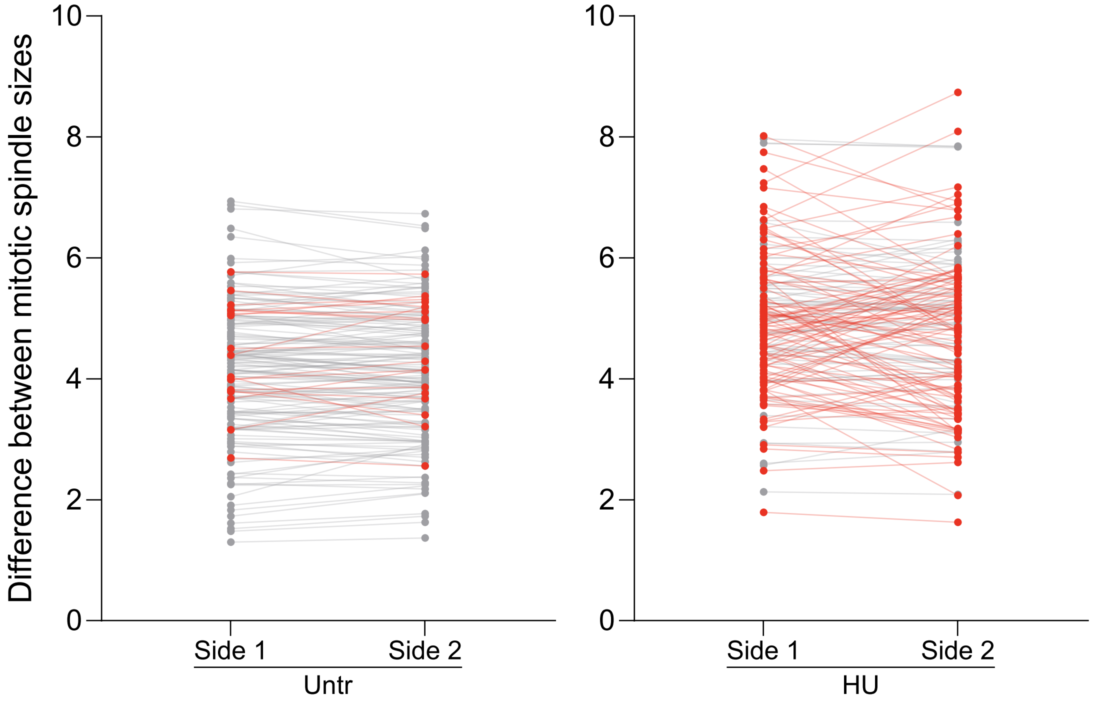
Difference between mitotic spindle sizes
10
8
6
4
2
0
Side 1
Side 2
Untr
10
8
6
4
2
0
Side 1
Side 2
HU

**Supplementary Figure 8: Detail of the mitotic spindle lengths in cells that were unperturbed or under replication stress.** Quantification of both sides of the spindle in unperturbed cells and mitosis after replicative stress, for each condition the lengths of the two sides are connected to show the low difference in unperturbed condition, and the high difference after replication stress, color scheme as in Figure 5 A and B.
